# TRPML1 signaling at lysosomes-mitochondria nexus drives triple-negative breast cancer mitophagy, metabolic reprogramming and chemoresistance

**DOI:** 10.1101/2025.09.11.675705

**Authors:** Alia K. Syeda, Shekoufeh Almasi, J. Cory Benson, Barry E. Kennedy, Ryan M. Yoast, Scott M. Emrich, Logan Slade, Shanmugasundaram Pakkiriswami, Vishnu V. Vijayan, Shashi Gujar, Thomas Pulinilkunnil, Mohamed Trebak, Yassine El Hiani

## Abstract

Inter-organelle signaling mechanisms, particularly those at the lysosomes-mitochondria interface, are critical for cancer cell metabolism, mitophagy and survival. However, the incomplete understanding of these mechanisms has limited the development of effective therapies, especially for triple-negative breast cancers (TNBC). Here, we demonstrate the lysosomal Ca²⁺-release channel TRPML1 as a master regulator of mitochondrial bioenergetics in TNBC cells. TRPML1 knockdown (ML1-KD) in TNBC cells selectively compromises mitochondrial respiration, reprograms cell metabolism, and induces mitochondrial fragmentation without impacting non-cancerous cells. Mitochondria of ML1-KD TNBC cells sequester around the endoplasmic reticulum (ER), increasing mitochondria-ER contact sites at the expense of mitochondria-lysosomes contacts. Mechanistically, ML1-KD reduces lysosomal acidification, thus hindering autophagic flux and completion of autophagy. ML1-KD inhibits TFEB-mediated mitophagy and oxidative defense mechanisms while causing mitochondrial Ca^2+^ overload, further impairing mitochondrial function. These alterations render ML1-KD TNBC cells highly sensitive to doxorubicin and paclitaxel at low doses that are typically ineffective on their own. Together, our findings establish TRPML1 as a critical inter-organelle regulator and highlight its potential as a therapeutic target to exploit the metabolic vulnerabilities of TNBC cells.

## INTRODUCTION

Cancer is characterized by uncontrolled cell growth and proliferation driven, at least in part, by perturbations in mitochondrial and lysosomal functions, two organelles central to tumor growth and survival (*1, 2*). Mitochondria are the energy powerhouse of the cell, essential in fueling biomass production and coordinating a variety of signaling pathways, including calcium (Ca²⁺) homeostasis, reactive oxygen species (ROS) balance, cell cycle regulation, and apoptosis—processes essential for cancer cell growth (*3–5*). Additionally, mitochondria-derived TCA cycle metabolites have recently emerged as important regulators of chromatin modifications, linking metabolic activity with gene expression in cancer cells (*6, 7*). Lysosomes, in turn, serve as a central signaling platform and metabolic regulator essential for cancer progression (*8, 9*). Beyond their canonical function of degrading and recycling damaged cellular components, lysosomes enable cancer cells to cope with nutrient deprivation and stress conditions by regulating key signaling pathways, including mammalian target of rapamycin (mTOR), AMP kinase (AMPK), and transcription factor EB (TFEB) (*10–13*). Together, these observations underscore that mitochondria and lysosomes act beyond their own boundaries to shape cellular signaling and determine cancer cell fate. As such, numerous studies have highlighted the fundamental role of the mitochondria-lysosomes interplay in maintaining cellular homeostasis and adapting to metabolic stress (*14, 15*). For example, the master transcriptional regulator of lysosomal function, TFEB has been shown to regulate mitophagy (*16, 17*), cell death and ATP production (*18, 19*). Disruption of lysosomal pH modulator V-ATPase was associated with diminished mitochondrial respiration and increased cell death (*20*). Lysosomal Ca²⁺ release has also been shown to influence mitochondrial Ca²⁺ homeostasis (*21*), thereby impacting critical mitochondrial functions such as Krebs cycle enzymatic activity and oxidative phosphorylation (*22*). Similarly, mitochondrial dysfunction is closely associated with impaired lysosomal activity. For instance, deletion of mitochondrial dynamics proteins (e.g. mitofusins, Dynamin-related protein 1) or chemical inhibition of the electron transport chain (ETC) impairs lysosomal activity, leading to the formation of large lysosomal vacuoles (*23, 24*). These bidirectional relationships suggest a functional interdependence between these two organelles, where the integrity of one organelle is closely tied to the other, and dysfunction in either compromises cellular homeostasis (*25, 26*). While this dynamic dependency supports cellular resilience, it also offers a metabolic vulnerability for cancer cells, which must balance rapid growth demands with nutrient availability and energy generation. Understanding the mechanisms that coordinate lysosome-mitochondria interaction could thus offer novel therapeutic opportunities for cancer. Defining the molecular mechanisms that coordinate lysosome–mitochondria communication may therefore uncover previously unrecognized vulnerabilities in cancer.

Recently, we (*27*) and others (*28*) have demonstrated the critical role of Transient Receptor Potential Mucolipin 1 (TRPML1), a lysosomal Ca²⁺-release channel, in promoting breast cancer survival and proliferation *in vitro* and in xenograft models. Specifically, we showed that downregulation of TRPML1 impairs TNBC growth and invasion by inhibiting mTORC1 activity and facilitating the release of lysosomal ATP into the extracellular space, respectively. Here, we define TRPML1 as a central, non-redundant regulator of lysosome–mitochondria communication in TNBC. Our findings reveal that TRPML1 downregulation alters mitochondrial structure and bioenergetics, disrupts mitophagy and autophagy flux, and remodels organellar contact sites, decreasing mitochondria-lysosomes contacts while increasing mitochondria-ER proximity. These alterations drive metabolic stress, cell cycle arrest, caspase-independent apoptosis, and enhanced chemosensitivity. Consistent with these cellular phenotypes, complementary analysis using the public OncoGenomics database [GSE22133−GPL5345 (*29*) and GSE-42568 (*30*)] supported our findings, showing that breast cancer patients with low TRPML1 mRNA expression have better clinical prognoses and higher survival rates. Collectively, these data position TRPML1 as a tractable therapeutic target in TNBC and underscore mechanistic non-redundancy with the related mucolipin TRPML3 (*31*), with TRPML1 operating in an acidic lysosomal pH window and through a signaling architecture distinct from TRPML3 (*31, 32*).

## RESULTS

### 1. TRPML1 knockdown induces G0/G1 cell cycle arrest and caspase-independent apoptosis in MDA-MB-231 cells

To determine the function of TRPML1 in the growth of triple-negative breast cancer cells MDA-MB-231, we examined the effects of TRPML1 stable knockdown using shRNA (ML1-KD) on colony formation. ML1-KD in MDA-MB-231 cells showed a significant reduction in the number of colonies compared to PLKO control cells (**Figure 1A**). In contrast, in the non-cancerous MCF10A cells, ML1-KD had no detectable effect on colony formation, highlighting the cancer cell-specific antiproliferative effect of TRPML1 downregulation. To further investigate the underlying mechanisms in MDA-MB-231 cells, we examined cell cycle distribution and apoptosis markers. ML1-KD induced cell cycle arrest at the G0/G1 phase (**Figure 1B**), as evidenced by decreased levels of Cyclin B1 and Cyclin D1, increased expression of p27, and reduced levels of E2F1 (**Figure 1C**), a critical transcription factor for cell cycle progression (*33*). Notably, these alterations in cell cycle markers were not observed in MCF10A cells (**Figure 1C**). ML1-KD also significantly increased the proportion of Annexin V-positive cells, indicating increased apoptosis (**Figure 1D**). Analysis of cell lysates revealed elevated levels of cytochrome C and cleaved caspase 9, but no detectable cleaved caspase 3 or 7, suggesting the involvement of a caspase-3/7 independent apoptotic pathway (**Figure 1E**). This was further supported by decreased phosphorylated BCL2 (pBCL2(S70)) and increased endonuclease G (EndoG) levels, whereas levels of apoptosis-inducing factor (AIF) remained unchanged (**Figure 1F**), suggesting a selective involvement of EndoG. Importantly, cytochrome C or EndoG levels were unaffected by ML1-KD in the non-cancerous MCF10A cells, reinforcing the selective effect of TRPML1 knockdown in TNBC. Together, these results indicate that TRPML1 knockdown inhibits proliferation of MDA-MB-231 by inducing G0/G1 cycle arrest and causing caspase-independent apoptosis, consistent with downstream mitochondrial dysfunction (*34*). These effects underscore a cancer-selective role of TRPML1 in controlling cell cycle progression and apoptosis.

**Figure 1:**
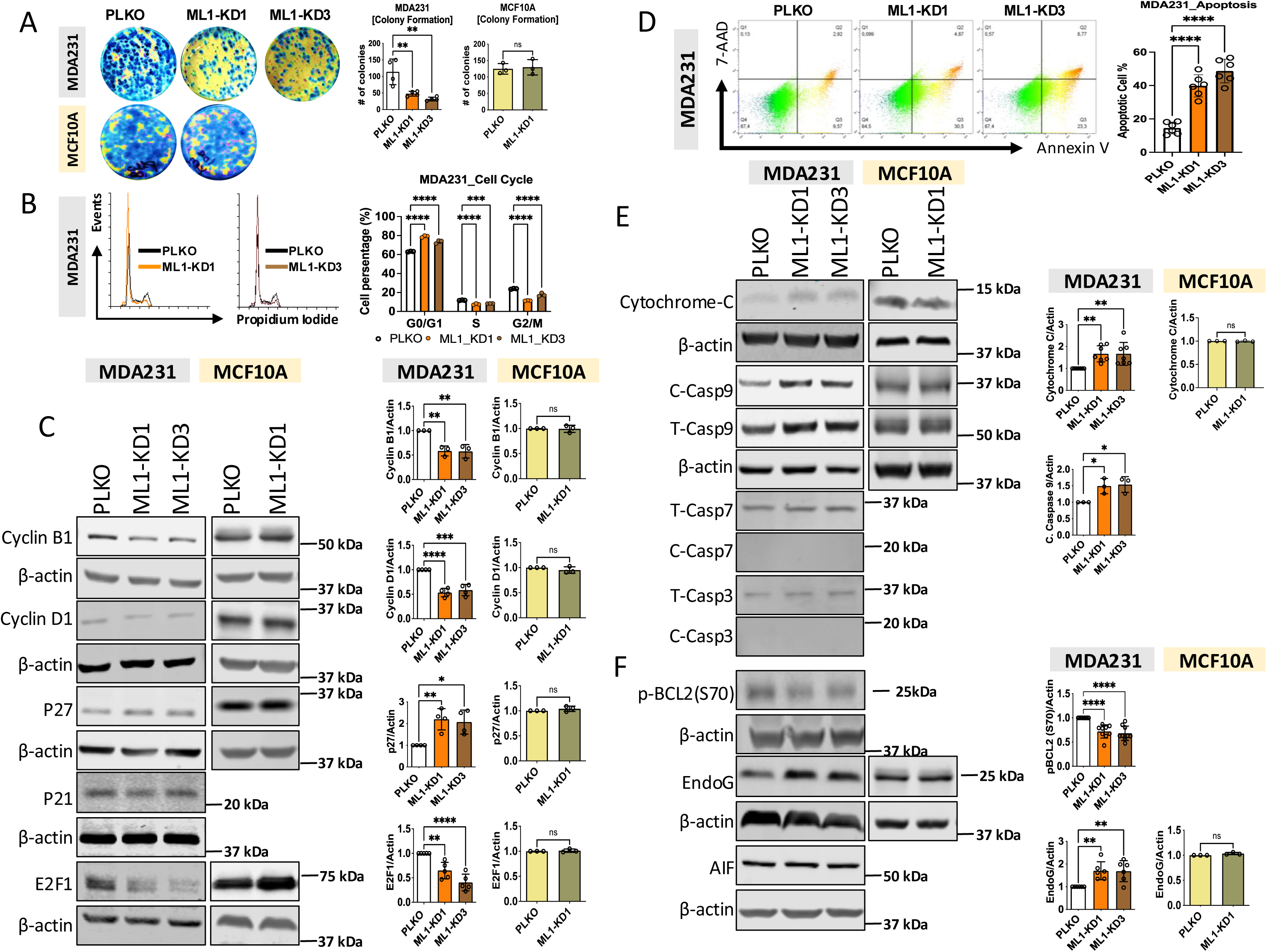
Downregulation of TRPML1 inhibits TNBC cell proliferation by inducing cell accumulation in G0/G1 phase of cell cycle and enhancing caspase-independent apoptosis. (A) Representative images of colony formation assay of PLKO and ML-KD in MDA-MB-231 and MCF10A cells; Graphs (Mean ± SD) show corresponding results from three independent experiments. (B) Representative flow plots gated on live cells for cell cycle analysis by fluorescence-activated cell sorting (FACS), using propidium iodide (PI) staining in MDA-MB-231 PLKO and ML1-KD conditions. Graphs (Mean ± SD) show corresponding percentages (Mean ± SD) of cell cycle phases (G0/1, S, G2/M), from a representative data out of three independent experiments; (C) Representative Western blot for Cyclin B1, Cyclin D1, P27, P21, and E2F1 in lysates from PLKO and ML1-KD of MDA-MB-231 and MCF10A cells – β-Actin served as a loading control. Graphs (Mean ± SD) show corresponding quantifications of protein levels normalized to β-Actin and reported to PLKO condition, from three to five independent experiments; (D) Representative flow plots gated on live cells for apoptosis analysis, using annexin V staining in MDA-MB-231 PLKO and ML1-KD conditions. Graph (Mean ± SD) shows percentages of apoptotic cells, from a representative data out of three independent experiments; (E and F) Representative Western blot for caspase-dependent apoptosis proteins Cytochrome C, caspase 9, caspase 7, and caspase 3 (E), and caspase-independent apoptosis proteins p-BCL2(S70), EndoG, and AIF (F) in lysates from PLKO and ML1 KD of MDA-MB-231 and MCF10A cells – β-Actin served as a loading control. Graphs (Mean ± SD) show corresponding quantifications of protein levels normalized to β-Actin and reported to PLKO condition, from three to nine independent experiments. Statistics: one-way ANOVA with Tukey’s multiple comparison test for PLKO vs ML1-KD1/ML1-KD3) of MDA-MB-231, and unpaired t-test and Mann-Whitney U test for PLKO and MLand unpaired t-test and Mann-Whitney U test for PLKO and ML1-KD1 of MCF10A cells. \**P* < 0.05, \*\**P* < 0.01, and \*\*\**P* < 0.001. ns, not significant.

### 2. TRPML1 knockdown impairs mitochondrial function and alters cellular bioenergetics of MDA-MB-231 cells

To determine whether the impaired proliferation and apoptosis observed in ML1-KD cells were associated with altered mitochondrial activity, we performed the Seahorse XF Mito Stress Test in MDA-MB-231 and MCF10A cells. Oxygen consumption rate (OCR) and extracellular acidification rate (ECAR) were recorded in real time following sequential injection of mitochondrial inhibitors **(Figure 2A, B)**, and the OCR/ECAR ratio was calculated at basal, maximal, and ATP-linked stages to assess the balance between oxidative phosphorylation and glycolytic flux. ML1-KD in MDA-MB-231 cells led to a marked decrease in both basal and maximal OCR/ECAR ratios, as well as in OCR/ECAR-linked ATP production, indicating impaired mitochondrial respiration and a disrupted bioenergetic profile **(Figure 2A, C-E)**. In contrast, ML1-KD had no effect on OCR/ECAR ratios in MCF10A cells **(Figure 2B–E)**, suggesting that TRPML1 is selectively required to sustain mitochondrial function in TNBC cells but not in non-tumorigenic epithelial cells. Consistently, cellular ATP levels were significantly reduced in ML1-KD MDA-MB-231 cells compared to vector-only (PLKO) controls, while ATP content in MCF10A cells remained unchanged **(Figure 2F)**, confirming the role of TRPML1 in maintaining energy production specifically in cancer cells. To further validate mitochondrial dysfunction, we performed high-resolution oxygraphy in MDA-MB-231 cells. ML1-KD cells exhibited a blunted respiratory response to stimulation with ADP, succinate, pyruvate, and malate compared to PLKO control cells, consistent with impaired oxidative phosphorylation (**Figure 2G**). Western blot analysis showed that ML1-KD decreased the expression of key electron transport chain (ETC) components, specifically complexes I, II, and IV (**Figure 2H, I)**, and was accompanied by increased levels of both total and mitochondrial ROS (**Figure 2K, L**), indicating the onset of mitochondrial dysfunction. Notably, MCF10A cells showed no significant changes in ETC levels **(Figure 2H, J)**, or total and mitochondrial ROS (**Figure 2K, L**) upon ML1-KD. Together, these findings demonstrate that TRPML1 is specifically required to preserve mitochondrial respiration, ATP production, and redox homeostasis in MDA-MB-231 TNBC cells. This establishes TRPML1 as a unique regulator of bioenergetics in cancer cells, independent of its role in non-tumorigenic epithelia.

**Figure 2:**
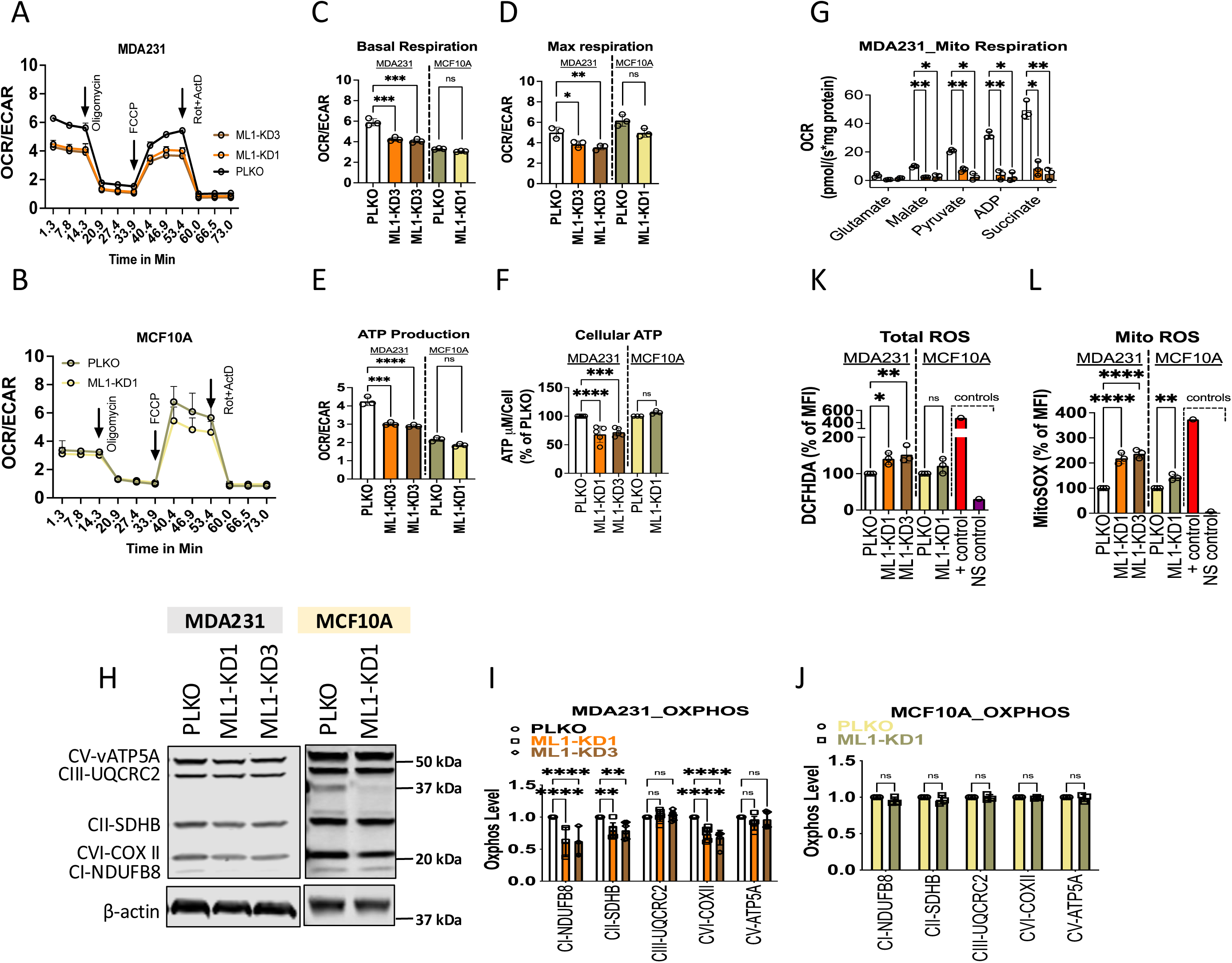
Downregulation of TRPML1 disturbs mitochondrial respiration, alters ECT protein levels, decrease ATP production and induces ROS and NO accumulation. (A and B) Overall OCR/ECAR ratio assessed by Seahorse analysis in PLKO and ML1-KD of MDA-MB-231 and MCF10A cells; Quantification of the Basal (C), maximal OCR/ECAR (D), and the OCR/ECAR linked-ATP (E). (F) Quantification of cellular ATP concentration using ATP Assay (Kit ab83355) in PLKO and ML1-KD of MDA-MB-231 and MCF10A cells. (G) Mitochondrial respiration OCR measured in PLKO and ML1-KD of MDA-MB-231, using Oroboros in response to treatment with glutamate, Malate, Pyruvate, ADP and Succinate. (H, I, J) Representative images (H) and quantification analysis of Western blot analysis of the subunits of OXPHOS complexes I–V in lysates from PLKO and ML1-KD of MDA-MB-231 (I) and MCF10A cells (J) – β-Actin served as a loading control. (K and L) Quantification of ROS using FACS for total ROS in stained cells with DCFHDA (K) and mitochondrial ROS for stained cells with MitoSOX (L), in PLKO and TRPML1 KD of MDA-MB-231 and MCF10A cells. All graphs (Means ± SD) are obtained from three independent experiments. \**P* < 0.05, \*\**P* < 0.01, and \*\*\**P* < 0.001. ns, not significant. Statistics: one-way ANOVA with Tukey’s multiple comparison test for PLKO vs ML1-KD1/ML1-KD3) of MDA-MB-231, and unpaired t-test and Mann-Whitney U test for PLKO and ML1-KD and unpaired t-test and Mann-Whitney U test for PLKO and ML1-KD1 of MCF10A cells. \**P* < 0.05, \*\**P* < 0.01, and \*\*\**P* < 0.001. ns, not significant.

### 3. TRPML1 knockdown induces mitochondrial fragmentation and alters organelle contacts in MDA-MB-231 cells

MitoTracker staining and confocal microscopy revealed that mitochondria in MDA-MB-231 PLKO control cells were uniformly distributed throughout the cytoplasm, whereas they appeared aggregated within the perinuclear region in ML1-KD MDA-MB-231 cells (**Figure 3A**). Quantitative analysis of confocal images revealed increased mitochondrial fragmentation and a significant reduction in mitochondrial volume in ML1-KD MDA-MB-231 cells compared to PLKO control cells, suggesting either increased mitochondrial fission or decreased fusion in ML1-KD cells (**Figure 3B**). Analysis of mitochondrial fission markers revealed no significant changes in mitochondrial fission factor (MFF) content and Ser616/Ser637 phosphorylation of Dynamin-related protein 1 (Drp1; **Figure 3C**). However, ML1-KD MDA-MB-231 cells showed a markedly decreased protein content of mitofusin 1 (MFN1), mitofusin 2 (MFN2), and optic atrophy 1 (OPA1), which are critical mediators of mitochondrial fusion (**Figure 3D**). These data indicate that TRPML1 knockdown tilts the balance in favor of mitochondrial fission and triggering the accumulation of fragmented mitochondria in MDA-MB 231 cells. Notably, we observed no changes in either mitochondrial structure or fusion protein levels in the non-cancerous MCF10A cells (**Figure 3A, E, F)**. Further, we analyzed mitochondrial morphology of ML1-KD MDA-MB-231 cells and their PLKO control counterparts using transmission electron microscopy (TEM). While we observed no differences in mitochondrial area between these two groups of cells (**Figure 4A, B**), ML1-KD MDA-MB-231 cells exhibited increased mitochondrial circularity and decreased aspect ratio compared to PLKO control cells (**Figure 4A, C, D**), both hallmark features of fragmented mitochondria (*35, 36*). Mitochondria of ML1-KD MDA-MB-231 cells also displayed increased cristae density compared to PLKO control cells (**Figure 4E**), potentially reflecting a compensatory adaptation to impaired mitochondrial respiration and reduced ATP production. Interestingly, TEM images and their quantification revealed that most mitochondria in ML1-KD MDA-MB-231 cells were surrounded by endomembranes, likely the ER (**Figure 4A, F**). These data indicate an increase in mitochondria-associated membranes (MAM) or mitochondria-ER contacts in ML1-KD MDA-MB-231 cells compared to PLKO control cells. Further, 3D Z-stack confocal microscopy showed a marked increase in the surface-to-surface contacts between MitoTracker-labeled mitochondria and Sec61-NEON-labeled ER (**Figure 4G, H**), indicating the spatial relationship between mitochondria and ER was altered following TRPML1 knockdown, with preferential shift towards enhanced ER-mitochondria proximity. Interestingly, we observed diminished co-localization of MitoTracker-stained mitochondria and LAMP1-labeled lysosomes (**Figure 4I, J**) in ML1-KD MDA-MB-231 cells compared to control cells. These data indicate that ML1-KD alters the balance of organellar contacts by bringing the ER closer to mitochondria while distancing them from the lysosomes, potentially altering the critical role lysosomes play in maintaining mitochondrial structure and function (*37*). Thus, TRPML1 loss uniquely remodels inter-organelle communication in TNBC cells, linking mitochondrial fragmentation with altered ER and lysosomal contacts.

**Figure 3:**
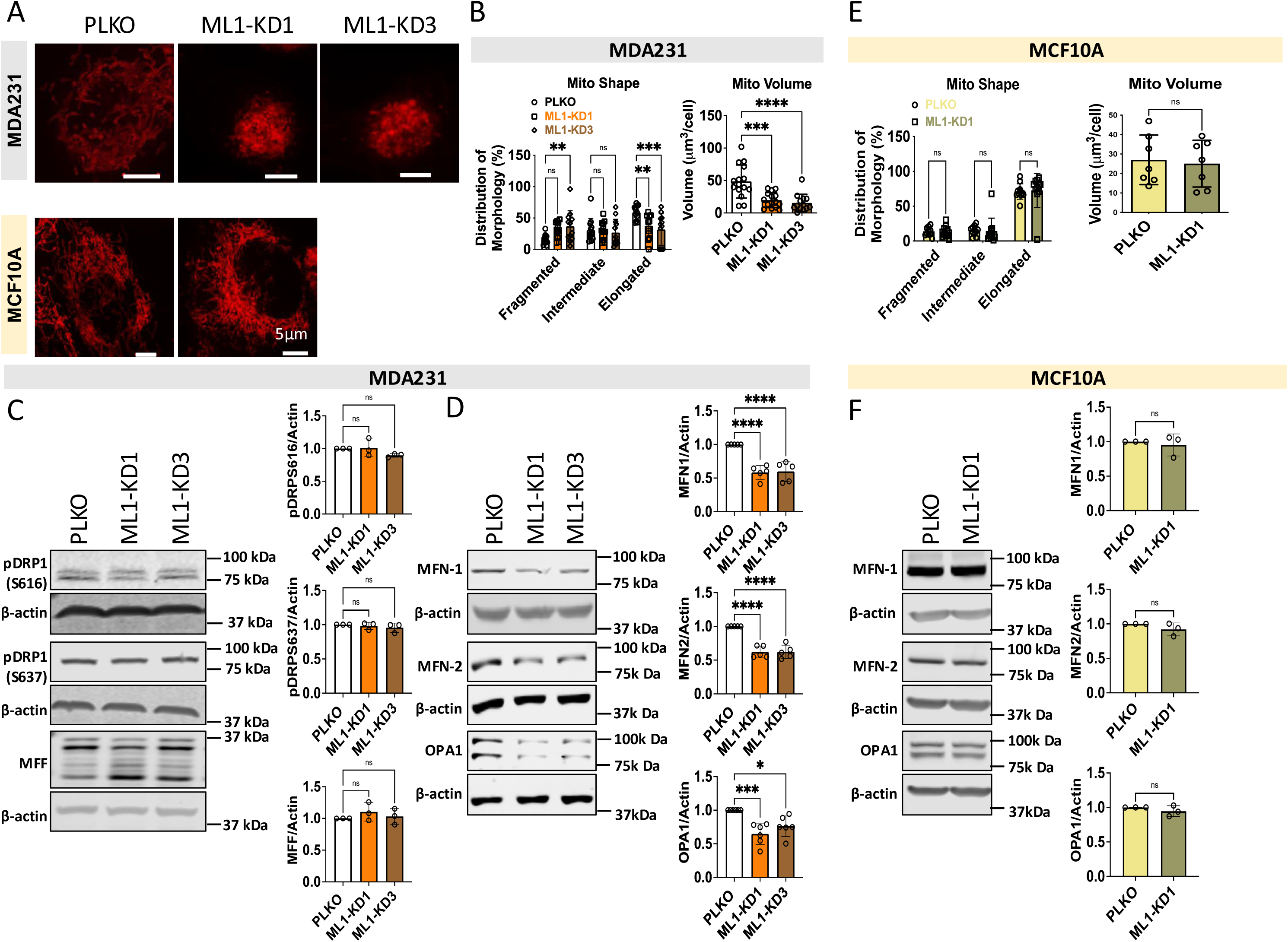
Downregulation of TRPML1 cause mitochondrial fragmentation, through the inhibition of mitochondrial fusion proteins. (A) Representative image of mitochondrial morphology assessed by MitotrackerRed^TM^ staining (maximum intensity projection shown), using confocal live-cell imaging using 100x magnification, in PLKO and ML1-KD from MDA-MB-231 and MCF10A – Scale bar: 5 um; (B and C) Graphs (Means ± SD) show quantitative analysis of mitochondrial length, using Imaris 10.2 software, distinguishing mitochondria into 3 main groups: small [fragmented]; longer [elongated]; average [intermediate] – in MDA-MB-231 (B) and MCF10A (E). Data were obtained from three independent biological experiments, 5-10 images per conditions, one cell per image; (C) Representative Western blot images and correspondent graphs for key mitochondrial fission (p-[s616]DRP1, p-[S637]DRP1 and MFF]) and (D) fusion (MFN1, MFN2 and OPA1) dynamic proteins in PLKO and ML1-KD of MDA-MB-231 cells; (F) Representative Western blot images and corresponding graphs for key mitochondrial fusion (MFN1, MFN2 and OPA1) in PLKO and MCF10A cells; Graphs (Mean ± SD) show quantifications of different protein levels normalized to β-Actin and reported to PLKO condition, from three to six independent experiments; Statistics: one-way ANOVA with Tukey’s multiple comparison test for PLKO vs ML1-KD1/ML1-KD3) of MDA-MB-231, and unpaired t-test and Mann-Whitney U test for PLKO and ML-1 KD and unpaired t-test and Mann-Whitney U test for PLKO and ML1-KD1 of MCF10A cells. \**P* < 0.05, \*\**P* < 0.01, and \*\*\**P* < 0.001. ns, not significant.

**Figure 4:**
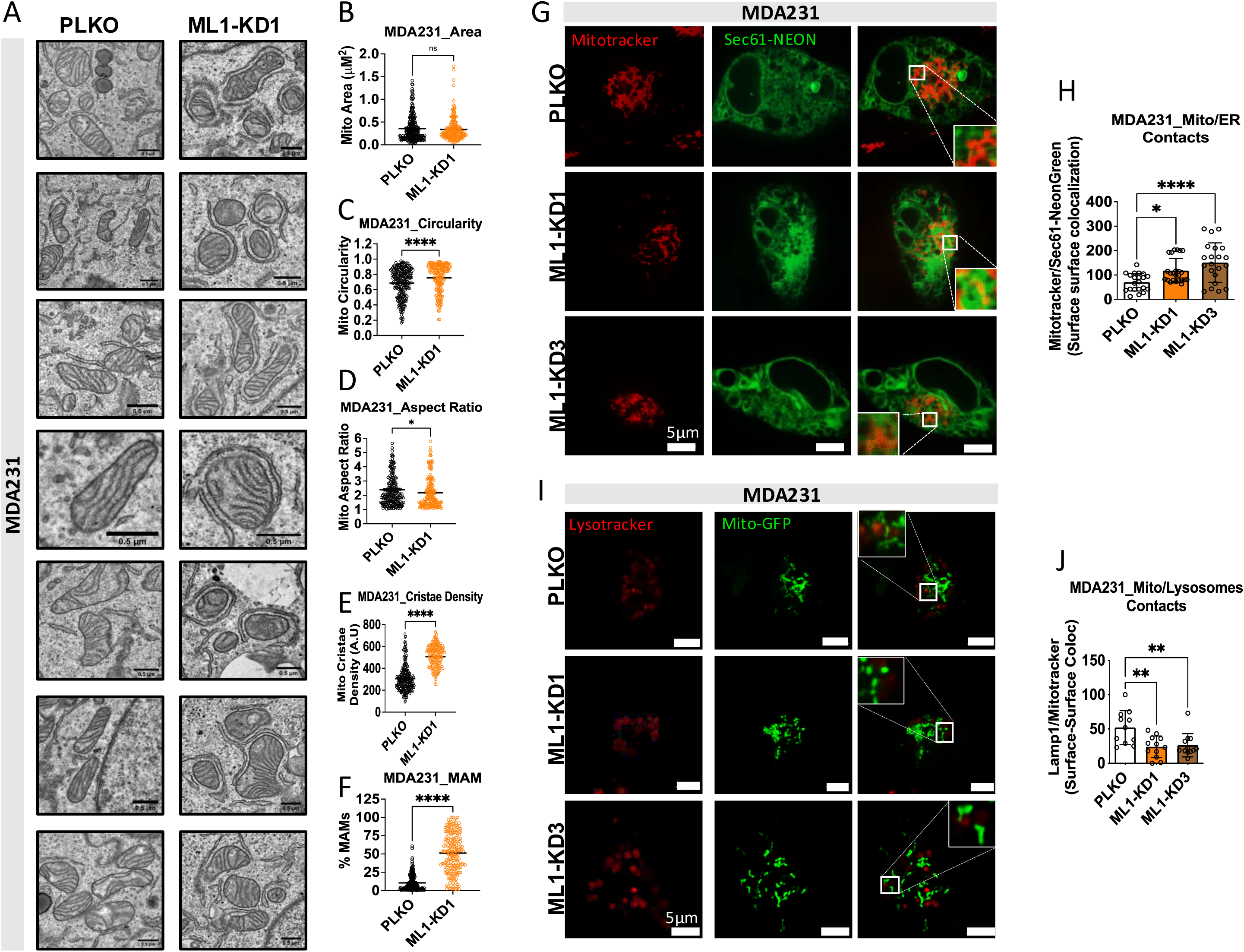
Downregulation of TRPML1 enhances contacts of mitochondria with ER while decreasing its contact with lysosomes. (A) Representative TEM images of PLKO and TRPML1 KD1 from MDA-MB-231 cells – Scale bar: 0.5 um. The area (B), circularity (C), Aspect Ratio (D), cristae density (E), and MAMs (F) of PLKO and TRPML1 KD1 in MDA-MB-231 cells were quantified using ImageJ. (G) Representative single Z-Stack planes, using confocal live-cell imaging with 100x magnification, from MDA-MB-231 PLKO and ML1-KD cells, stained with mitochondria marker (Mitotracker, red signal) and ER marker (Sec61-NEON, green signal) cells – Scale bar: 5 um; Merge panels show the degree of volume overlapping between mitochondria and the ER. (H) Graph (Means ± SD) represents the % of surface-surface colocalization between ER volume that are shared with mitochondrial volume in PLKO vs ML1-KD cells. Data were obtained from three independent biological experiments, 5-10 images per conditions, one cell per image; (I) Representative single Z-Stack planes, using confocal live-cell imaging with 100x magnification, from MDA-MB-231 PLKO and ML1-KD cells, stained with lysosome marker (Lysotracker, red signal) and mitochondria marker (Mitotracker, green signal) – Scale bar: 5 um; Merge panels show the degree of volume overlapping between lysosomes and mitochondria. (J) Graph (Means ± SD) represents the % of surface-surface colocalization between lysosomal volume that are shared with mitochondrial volume in PLKO vs ML1-KD cells – Data were obtained from three independent biological experiments, 5-10 images per conditions, one cell per image. Statistics: one-way ANOVA with Tukey’s multiple comparison test for PLKO vs ML1-KD1/ML1-KD3) of MDA-MB-231, and unpaired t-test. \**P* < 0.05, \*\**P* < 0.01, and \*\*\**P* < 0.001. ns, not significant.

### 4. TRPML1 knockdown disrupts autophagy and alters lysosomal fusion

TRPML1-mediated lysosomal Ca^2+^ release is critical for the activation of the mTOR pathway (*38*). mTORC1 phosphorylates P70S6K at Thr389 and ULK1 at Ser757, preventing autophagy (*39*). Furthermore, it is well known that increased AMP/ATP ratio leads to AMPK phosphorylation at Thr172 causing its activation and the subsequent phosphorylation of ULK1 at Ser555 by AMPK promote initiation of autophagy (*40*). Autophagy initiation is followed by complex signaling pathways leading to autophagosome formation and maturation, and culminating in autophagic flux, the process of fusion of autophagosomes with lysosomes and subsequent degradation of the autophagic cargo (*41–43*). Consistent with our previous findings, phosphorylation of P70S6K at Thr389 was decreased in ML1-KD MDA-MB-231 cells compared to PLKO control cells (**Figure 5A**), indicating suppression of mTOR activity. ML1-KD lysates also displayed reduced phosphorylation of ULK1 at Ser757, suggesting enhanced autophagy initiation (**Figure 5A**). In parallel, ML1-KD increased AMPK phosphorylation at Thr172 (**Figure 5B**) and increased phosphorylation of ULK1 at Ser555 (**Figure 5B**), further supporting increased autophagy initiation in response to TRPML1 knockdown (*44, 45*).

**Figure 5:**
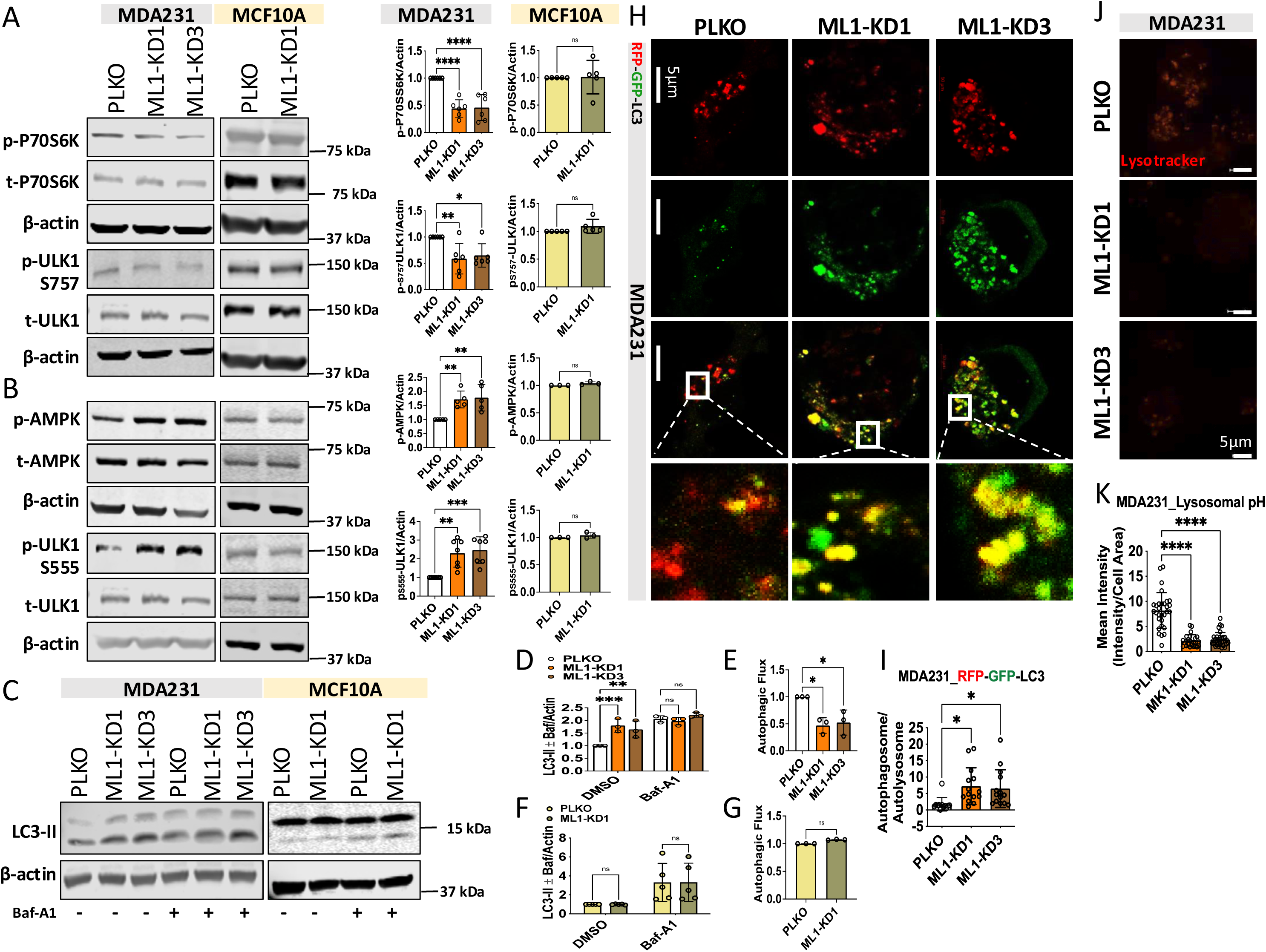
Downregulation of TRPML1 induces autophagy initiation but blocks autophagy flux. (A and B) Representative Western blot of mTOR and autophagy initiation markers, including phospho and total P70S6K (mTOR marker and target), phospho and total 757ULK1 (autophagy initiation marker, mTOR target), phospho and total 555ULK1 (autophagy initiation marker, AMPK target) in lysates from PLKO and ML1-KD of MDA-MB-231 and MCF10A cells; Graphs (Mean ± SD) show corresponding quantifications of protein levels normalized to β-Actin and reported to PLKO condition, from three to seven independent experiments. (C) Representative Western blot analysis of LC3-II in the presence or absence of bafilomycin A1 (BafA) in MDA-MB-231 and MCF10A PLKO and ML1-KD conditions – Cells were treated with DMSO or 200 nM BafA for 2 h, then cells were collected, lysed, and subjected to Western blot analysis. (D, F) Graph (Mean ± SD) shows quantification of LC3 II levels in PLKO vs ML1-KD in DMSO and BafA in MDA-MB-231 (D) and MCF10A (F) cells. Quantifications of LC3II protein levels was normalized to β-Actin and reported to PLKO condition, from three to five independent experiments; (E, G) The calculated autophagy flux: ratio of LC3II/ β-Actin in presence of BafA minus ration of LC3II/ β-Actin in absence of LC3II for each condition, normalized to PLKO condition. (H, I) Representative fluorescence images of MDA-MB-231 cells transiently expressing tandem RFP-GFP-LC3 to monitor autophagosome–lysosome fusion in PLKO and ML1-KD conditions, using confocal live-cell imaging with 63x magnification – Scale bar: 5 μm. Individual GFP and RFP channels, as well as merged images are shown. Quantification was performed using merged images, and the accompanying graph (Mean ± SD) displays the ratio of yellow puncta (GFP+RFP+; non-acidified autophagosomes, indicating blocked fusion) to red-only puncta (RFP+GFP−; acidified autolysosomes, indicating successful fusion) – Data were obtained from three independent biological experiments, 5-10 images per conditions, two to three cells per image ; (J, K) Representative fluorescence images of MDA-MB-231 stained with Lysotracker to monitor lysosomal pH in PLKO ML1-KD conditions, using confocal live-cell imaging with 40x magnification. Graphs (Mean ± SD) show corresponding quantifications of Lysotracker mean intensity per cell between PLKO and ML1-KD cells – Data were obtained from three independent biological experiments, 5-10 images per conditions, three to five cells per image. Statistics: one-way ANOVA with Tukey’s multiple comparison test for PLKO vs ML1-KD1/ML1-KD3) of MDA-MB-231, and unpaired t-test and Mann-Whitney U test for PLKO and ML-1 KD and unpaired t-test and Mann-Whitney U test for PLKO and ML1-KD1 of MCF10A cells. \**P* < 0.05, \*\**P* < 0.01, and \*\*\**P* < 0.001. ns, not significant.

To determine the effect of TRPML1 knockdown on the autophagic flux, we analyzed LC3-II lipidation levels in the presence and absence of bafilomycin A1 (BafA), an inhibitor of autophagosome-lysosome fusion (*46*). LC3-II which resides on autophagosomal membranes, serves as a marker of autophagosome abundance. Basal LC3-II levels were elevated in ML1-KD compared to PLKO control cells in BafA untreated MDA-MB-23 cells **(Figure 5C, D)**, suggesting either enhanced autophagosome formation or impaired degradation. However, BafA treatment failed to further increase LC3-II levels in ML1-KD compared PLKO cells **(Figure 5C, D)**, indicating that the autophagosomes of ML1-KD were already unable to fuse with lysosomes, even in the absence of BafA. This result implies that the accumulation of LC3-II in ML1-KD cells results from impaired autophagic flux, not increased autophagosome production **(Figure 5E)**. Notably, TRPML1 knockdown did not affect autophagy markers or flux in non-cancerous MCF10A cells (**Figure 5A-C, F, G)**). To further corroborate these findings, we monitored autophagosome-lysosome fusion by transfecting cells with a reporter plasmid pMRX-IP-GFP-LC3-RFP expressing a tandem of pH sensitive EGFP with pH resistant mRFP1 (*47*). In this assay, autophagosomes exhibit yellow puncta due to colocalization of GFP (green) with RFP (red). However, upon fusion with lysosomes, the acidic environment quenches the GFP, leaving only the RFP red signal in autolysosomes, indicative of successful autophagosome-lysosome fusion and lysosomal proteolytic action (*48*). MDA-M-231 PLKO control cells showed increased red fluorescence, consistent with elevated autophagic flux. However, ML1-KD MDA-MB-231 cells displayed predominantly yellow puncta, confirming a block in autophagosome-lysosome fusion (**Figure 5H, I**).

Given that lysosomal acidity is essential for fusion and degradation (*49–52*), we next assessed lysosomal pH, using LysoTracker staining as an indirect measure of lysosomal acidification (*31*). ML1-KD MDA-MB-231 cells displayed reduced LysoTracker intensity compared to PLKO control cells, indicating lysosomal alkalinization (**Figure 5J, K**). This finding supports the conclusion that impaired autophagosome-lysosome fusion in ML1-KD cells is associated with compromised lysosomal acidification. Taken together, these results demonstrate that TRPML1 knockdown in MDA-MB-231 cells exerts a dual effect: 1) it promotes autophagy initiation by inhibiting mTOR signaling, and 2) it disrupts the autophagic flux by inducing lysosomal alkalinization and blocking autophagosome-lysosome fusion. This identifies a unique cancer-specific defect in autophagy regulation downstream of TRPML1, underscoring its role in maintaining lysosome–mitochondria homeostasis.

### 5. Knockdown of TRPML1 Suppresses Mitophagy via TFEB-Mediated Parkin/Pink1 Pathways

We next investigated whether the impaired lysosomal function in ML1-KD MDA-MB-231 cells affects mitophagy and mitochondrial protein quality control, which could explain the quiescent metabolic profile observed in these cells. To evaluate mitophagy, we first examined the spatial relationship between mitochondria (labelled with green Mito-GFP) and LC3-positive autophagic vesicles (marked by red LC3-mCherry) (**Figure 6A**), as the proximity of mitochondria to LC3-positive structure is a critical step in mitophagy (*53*). ML1-KD MDA-MB-231 cells showed less contact between mitochondria and LC3-labeled vesicles compared to PLKO controls (**Figure 6B**), suggesting impaired mitophagosome formation and reduced targeting of mitochondria for degradation. These findings are consistent with our earlier data showing that mitochondria are more distant from lysosomes in ML1-KD MDA-MB-231 cells (**Figure 4I, J**). Quantification of LC3 puncta further revealed an increased number in ML1-KD MDA-MB-231 cells (**Figure 6C**), suggesting excessive accumulation of autophagosomes consistent with altered mitophagy.

**Figure 6:**
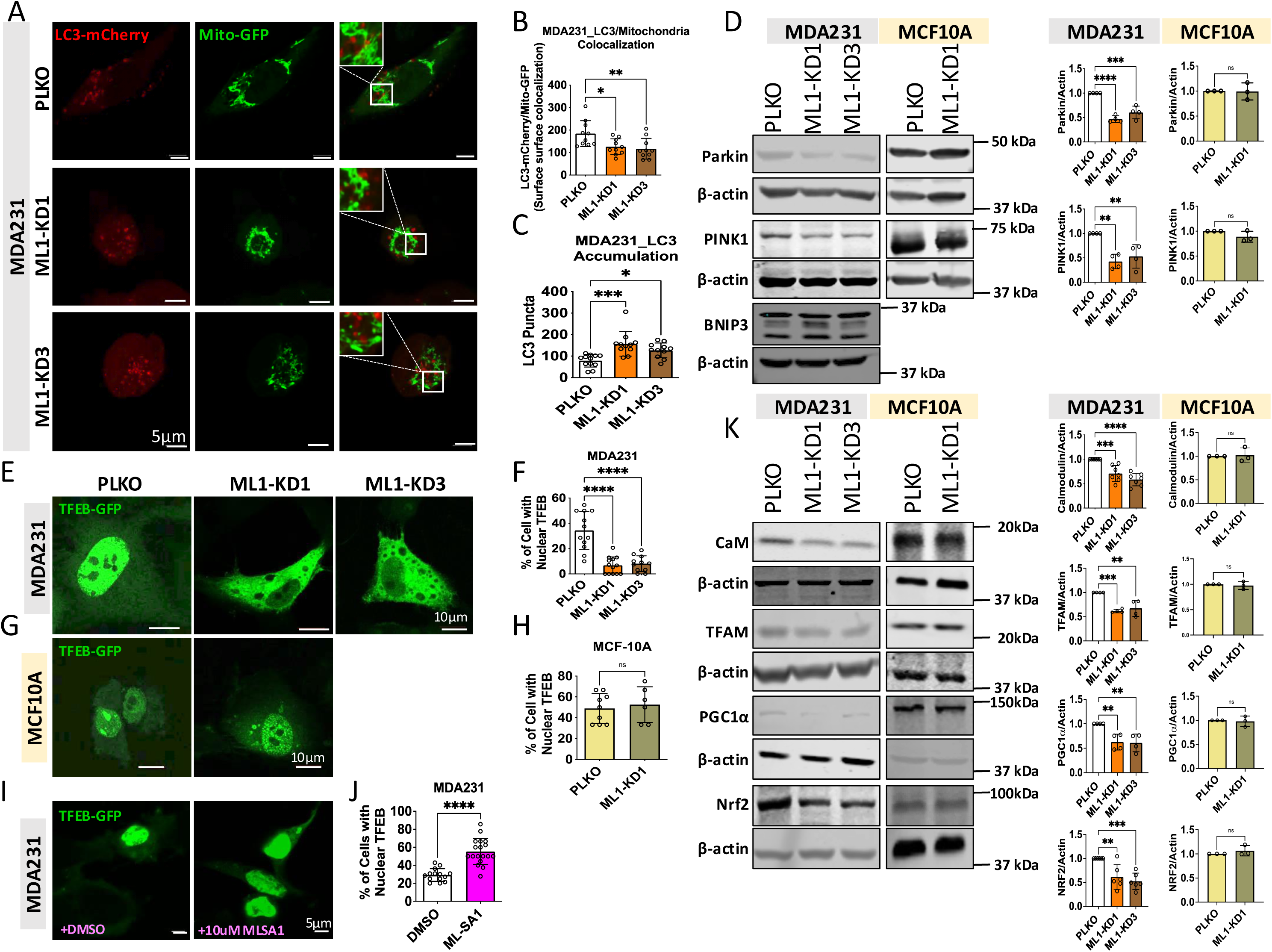
Downregulation of TRPML1 alters mitophagy through inhibition of CaM/TFEB pathway. (A) Representative single Z-Stack planes, using confocal live-cell imaging with 100x magnification, from MDA-MB-231 PLKO and ML1-KD cells, co-transfected with LC3-mCherry and Mito-GFP to assess mitophagy. Red (LC3) and green (mitochondria) channels, as well as merged images, are shown – Scale bar: 5 um. Merge panels show the degree of volume overlapping between autolysosomes and mitochondria. (B) Graph (Means ± SD) represents the % of surface-surface colocalization between autolysosomal volume that are shared with mitochondrial volume in PLKO vs ML1-KD cells. (C) LC3-mCherry puncta (autolysosomes) were quantified to evaluate LC3 levels in PLKO vs ML1-KD conditions Data were obtained from three independent biological experiments, 5-10 images per conditions, one cell per image; (D) Representative Western blot of mitophagy markers, including Parlin, Pink1, and BNIP3 from PLKO and ML1 KD of MDA-MB-231 and MCF10A cells. Graphs (Mean ± SD) show corresponding quantifications of protein levels normalized to β-Actin and reported to PLKO condition, from three to four independent experiments. (E) Representative images of MDA-MB-231 cells transfected with TFEB-GFP in PLKO and ML1-KD conditions, using confocal live-cell imaging with 40x magnification – Scale bar: 5 um. (F) Graph (Mean ± SD) shows corresponding quantifications as % of cell with nuclear TFEB in total cells per field, 5-10 images per conditions, five to 10 cells per image. (G) Representative images of MCF10A cells transfected with TFEB-GFP in PLKO and ML1-KD conditions, using confocal live-cell imaging with 40x magnification – Scale bar: 5 um. (H) Graph (Mean ± SD) shows corresponding quantifications as % of cell with nuclear TFEB in total cells per field, 5-10 images per conditions, three to seven cells per image. (I, J) Representative images of MDA-MB-231 cells transfected with TFEB-GFP in presence and absence of 10uM ML-SA1, ML1 activator, using confocal live-cell imaging with 40x magnification – Scale bar: 5 um. Graph (Mean ± SD) shows corresponding quantifications as % of cell with nuclear TFEB in total cells per field, 5-10 images per conditions, five to 10 cells per image; (K) Representative Western blot of TFEB upstream marker (calmodulin) and TFEB downstream markers (TFAM, PCG1a, and NRF2) from PLKO and ML1 KD of MDA-MB-231 and MCF10A cells. Graphs (Mean ± SD) show corresponding quantifications of protein levels normalized to β-Actin and reported to PLKO condition, from three to seven independent experiments. Statistics: one-way ANOVA with Tukey’s multiple comparison test for PLKO vs ML1-KD1/ML1-KD3) of MDA-MB-231, and unpaired t-test and Mann-Whitney U test for PLKO and ML-1 KD and unpaired t-test and Mann-Whitney U test for PLKO and ML1-KD1 of MCF10A cells. *P < 0.05, **P < 0.01, and ***P < 0.001. ns, not significant.

To determine the mechanisms behind impaired mitophagy, we examined the expression levels of key mitophagy regulators such as Parkin, Pink1, and BNIP3 (*54, 55*). ML1-KD MDA-MB-231 cells exhibited markedly reduced expression of Pink1 and Parkin, but not BNIP3 (**Figure 6D**), pointing to suppression of canonical PINK1-Parkin pathway. Consequently, we examined ML1-KD cells for potential changes in TFEB activation, a known upstream regulator of Pink1 and Parkin (*56, 57*). TFEB nuclear localization was significantly diminished in ML1-KD MDA-MB-231 cells compared to PLKO controls (**Figure 6E, F**). In contrast, pharmacological activation of the TRPML1 using ML1-SA1 (10 μM) enhanced TFEB nuclear translocation in MDA-MB-231 PLKO cells (**Figure 6I, J**), consistent with previous reports indicating that TRPML1 positively regulates TFEB activity (*58*). We further investigated potential upstream regulators and downstream effectors of TFEB (*59*). Calmodulin, a Ca^2+^-binding protein that activates calcineurin, a phosphatase that dephosphorylates TFEB leading to its activation and nuclear translocation (*58*), was decreased in ML1-KD MDA-MB-231 cells **(Figure 6K**). In addition, TFEB downstream responsive proteins, including PGC-1α (key regulator of mitochondrial biogenesis (*60*)), TFAM (essential for mitochondrial DNA maintenance and transcription (*61*)), and NRF2 (involved in oxidative defense (*62*)) were also decreased, indicating the inhibition of TFEB-mediated programs linked to mitophagy (**Figure 6K**). Importantly, TRPML1 knockdown had no effect on mitophagy markers or TFEB translocation in non-cancerous MCF10A cells (**Figure 6D-K**), highlighting the specific effects of TRPML1 in MDA-MB-231 TNBC cells. Together, these findings suggest that TRPML1 knockdown disrupts mitophagy in MDA-MB-231 cells by inhibiting lysosomal Ca^2+^ release, which in turn blocks TFEB nuclear translocation and suppresses the induction of essential proteins for mitochondrial maintenance, biogenesis and redox balance. This identifies a cancer-selective defect in mitophagy control downstream of TRPML1 and underscores its role in coordinating lysosomal signaling with mitochondrial quality control.

### 6. TRPML1 knockdown alters cytosolic and mitochondrial Ca^2+^ signaling

TRPML1-mediated lysosomal Ca^2+^ signaling regulates a plethora of cellular processes including exocytosis, phagocytosis, autophagy and Ca^2+^ shuttling to the ER and mitochondria (*21, 63*). Specifically, lysosomal Ca^2+^ release by TRPML1 facilitates Ca^2+^ transfer to mitochondria, a process dependent on mitochondria-lysosome contact sites (*21*). Loss of TRPML1 function, as seen in the lysosomal storage disorder mucolipidosis type IV (MLIV), disrupts these contacts and impairs contact-dependent mitochondrial Ca^2+^ uptake (*21*). In light of the mitochondrial defects observed in ML1-KD MDA-MB-231 cells, we investigated whether TRPML1-mediated lysosomal Ca^2+^ release directly contributes to mitochondrial Ca^2+^ uptake (**Figure 7A**), a key determinant of mitochondrial quality control and metabolic activity (*3, 64*). To this end, we activated TRPML1 using the agonist ML1-SA1 and monitored mitochondrial Ca^2+^ uptake using Rhod-2 dye, both in the absence and presence of extracellular Ca^2+^. Notably, 50 µM of TRPML1-SA1 (a high concentration) failed to induce any mitochondrial Ca^2+^ uptake in either MDA-MB-231 or MCF10A cells when extracellular Ca^2+^ was absent, regardless of TRPML1 expression status (ML1-KD and PLKO cells) (**Figure 7B**). Even when TRPML1 was overexpressed in MDA-MB-231 cells, mitochondrial Ca^2+^ levels remained undetectable under free extracellular Ca^2+^, suggesting that TRPML1 does not directly mediate lysosome-to-mitochondria Ca^2+^ transfer (**Figure 7C**), despite prior reports of contact-dependent transfer (*21*). In contrast, when extracellular Ca^2+^ was present, ML1-SA1 led to a dose-dependent increase in mitochondrial Ca^2+^ uptake in PLKO control MDA-MB-231 cells that was particularly evident at 25-50 μM of ML1-SA1 (**Figure 7D_1_, D_4_, D_5_**). Strikingly, mitochondrial Ca^2+^ uptake was significantly enhanced in ML1-KD MDA-MB-231 cells and manifested at the lower ML1-SA1 concentration of 5 μM (**Figure 7D_2_-D_5_**).

**Figure 7.**
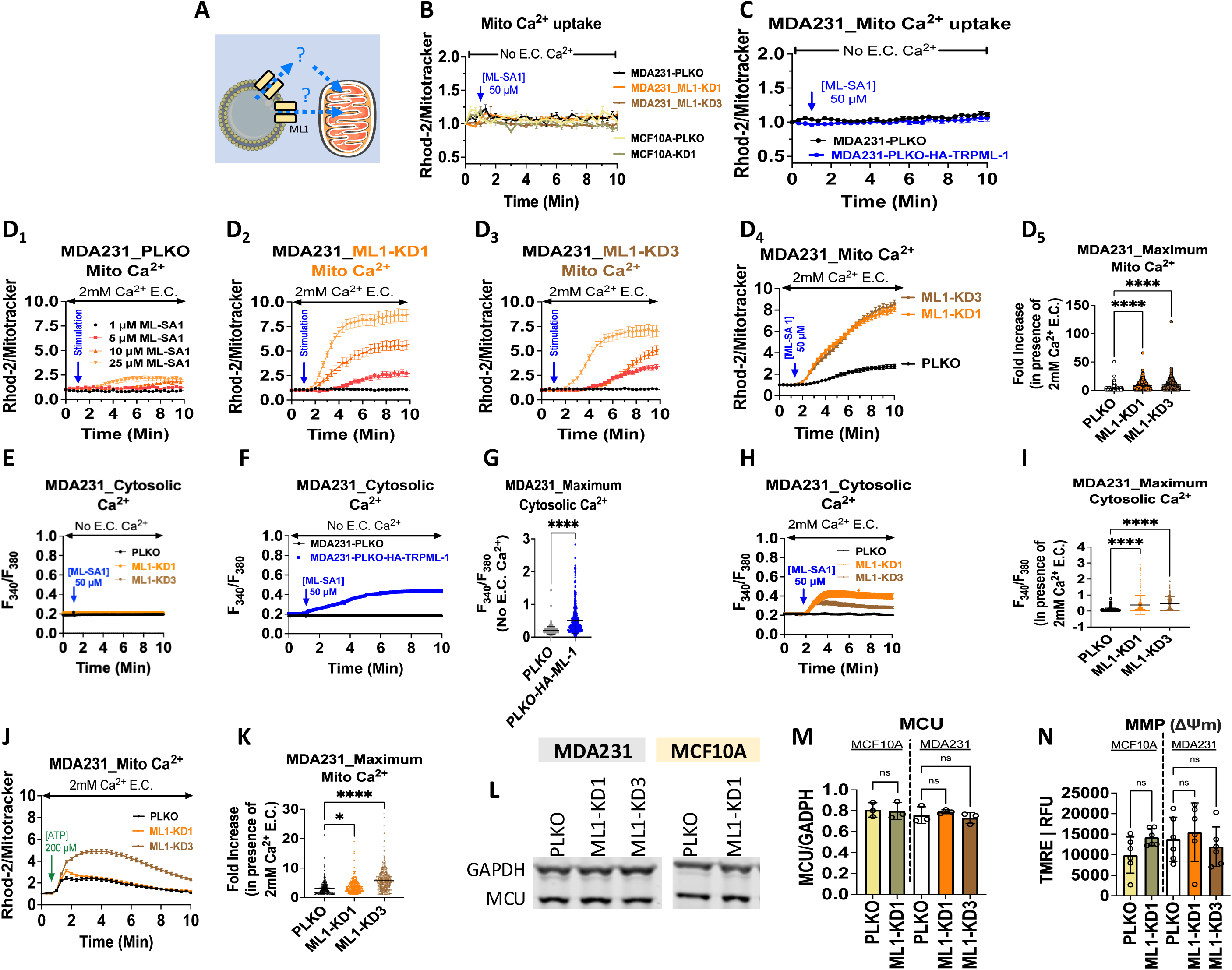
Downregulation of TRPML1 indirectly induces mitochondrial Ca^2+^ uptake. (A) schematic presentation highlighting the question whether lysosomal TRPML1 activation can directly influence mitochondrial Ca^2+^ uptake or via an intermediary pathway. (B) Traces representing mitochondrial Ca^2+^ uptake in MDA-MB-231 PLKOs, ML-1KD1/ML1-KD3 and MCF-10A in response to ML1-SA1 stimulation in the absence of extracellular Ca^2+^. Traces represent the mean ± SEM. (C) Traces representing mitochondrial Ca^2+^ uptake in MDA-MB-231 PLKOs and PLKOs transfected with HA-TRPML1 in response to ML1-SA1 in the absence of extracellular Ca^2+^. Traces represent the mean ± SEM (D_1-4_) Traces representing mitochondrial Ca^2+^ uptake in response to increasing doses of ML1-SA1 in MDA-MB-231 PLKOs and ML1-KD1/ML1-KD3s. (D_5_) Quantification of maximum mitochondrial Ca^2+^uptake in (D_4_), error bars represent the mean ± SD, results analyzed by Kruskal-Wallis test with Dunn’s multiple comparison correction. (E) Traces representing cytosolic Ca^2+^ uptake in MDA-MB-231 PLKOs vs ML1-KD1/ML1-KD3 in response to ML1-SA1 stimulation in the absence of extracellular Ca^2+^. Traces represent mean ± SEM. (F) Traces representing cytosolic Ca^2+^ uptake in MDA-MB-231 PLKOs vs PLKOs transfected with HA-TRPML-1 in response to ML1-SA1 in the absence of extracellular Ca^2+^. Traces represent mean ± SEM. (G) Quantification of (F), results analyzed by Mann Whitney test. (H) Traces representing cytosolic Ca^2+^ uptake in MDA-MB-231 PLKOs vs ML1-KD1/ML1-KD3 cells in response to ML1-SA1 stimulation in the presence of extracellular Ca^2+^. Traces represent the mean ± SEM. (I) Quantification of (H), error bars represent the mean ± SD, results analyzed by Kruskal Wallis test with post-hoc Dunn’s multiple comparisons test. (J) Traces to represent mitochondrial Ca^2+^ uptake in MDA-MB-231 PLKO vs ML1-KD1/ML1-KD3 cells in response to stimulation with 200 µM ATP. (K) Corresponding quantification of (J), error bars are representative of mean ± SD, results analyzed by Kruskal Wallis test with post-hoc Dunn’s multiple comparisons test. (L) Representative western blot images measuring MCU expression in MDA-MB-231 PLKO, ML-1KD1/ML-1-KD3 and MCF-10A. (M) Quantification of (L) with MCU protein expression normalized to GAPDH loading control, graph represents means ± SD. Results analyzed by one-way ANOVA with post-hoc Tukey’s multiple comparisons test. (N) Quantification of TMRE measurements of MDA-MB-231 PLKO, ML1-KD1/ML1-KD3, and MCF-10A. Results analyzed by one-way ANOVA with post-hoc Tukey’s multiple comparison test. All mitochondrial Ca^2+^ measurements were obtained with Rhod-2 AM and normalized to MitoTracker^TM^ Green. All cytosolic Ca^2+^ measurements were obtained with Fura-2 AM. All presented experiments represent three biological replicates. *P < 0.05, **P < 0.01, ***P < 0.001 and ****P <0.0001, ns, not significant.

We also tested whether the activation of TRPML1 with TRPML1-SA1 could activate Ca^2+^ release into the cytosol by monitoring cytosolic Ca^2+^ levels with Fura2 in the absence of extracellular Ca^2+^ (**Figure 7 E-G**). Remarkably, TRPML1-SA1 at the high concentration of 50 µM did not affect cytosolic Ca^2+^ levels in both PLKO control and ML1-KD MDA-MB-231 cells when extracellular Ca^2+^ was omitted (**Figure 7 E)**. However, when TRPML1 was overexpressed in PLKO MDA-MB-231 cells, a significant increase in cytosolic Ca^2+^ was detected (**Figure 7 F, G)**, suggesting that endogenous TRPML1 activity is minimal under baseline conditions/protocols but become apparent upon overexpression. Paradoxically, in the presence of extracellular Ca^2+^, ML1-SA1 caused a greater increase in cytosolic Ca^2+^ in ML1-KD MDA-MB-231 cells compared to PLKO control cells (**Figure 7 H, I)**. These data suggest that ML1-SA1 has off-target effects, potentially activating plasma membrane Ca^2+^ influx pathways that are upregulated or sensitized in the absence of TRPML1. Regardless of the specificity of TRPML1-SA1, our results do not support a direct lysosomes-to-mitochondria Ca^2+^ transfer mediated by TRPML1. Rather, the increase in mitochondrial Ca^2+^ uptake in ML1-KD cells appears to result from elevated cytosolic Ca^2+^, likely amplified by ER Ca^2+^ release. Supporting this, receptor stimulation with ATP under physiological conditions led to significantly greater mitochondrial Ca^2+^ uptake in ML1-KD MDA-MB-231 cells compared to PLKO control cells (**Figure 7 J, K)**. The increased mitochondrial Ca^2+^ uptake in ML1-KD MDA-MB-231 cells was neither due to increased mitochondrial Ca^2+^ uniporter (MCU) protein levels (**Figure 7L, M)**, nor to enhanced mitochondrial driving force through changes in mitochondrial membrane potential (**Figure 7N)**. Therefore, we propose that elevated mitochondrial Ca^2+^ uptake observed in ML1-KD MDA-MB-231 cells is likely driven by increased mitochondria-ER contact sites we have uncovered in these cells (**Figure 4A, F, G, H**). Together, these data position TRPML1 as a regulator of Ca²⁺ microdomain coupling and organelle contact remodeling in TNBC cells, linking cytosolic/ER Ca²⁺ dynamics to mitochondrial Ca²⁺ loading without invoking direct lysosome-to-mitochondria transfer.

### 7. TRPML1 knockdown remodels the metabolism of MDA-MB-231 cells

Given the altered mitochondrial architecture, disrupted bioenergetics, impaired Ca^2+^ homeostasis, and remodeled organellar contact sites observed in ML1-KD MDA-MB-231 cells, we sought to determine whether these defects were associated with a distinct mitochondrial metabolic signature. To address this, we performed targeted mass spectrometry-based metabolomics on both PLKO and ML1-KD MDA-MB-231 and MCF10A cells. Consistent with the non-essential role of TRPML1 in MCF10A cells, ML1-KD induced only minimal changes in the overall metabolomic profile of MCF10A cells compared to PLKO cells (**Figure 8 A**). In striking contrast, TRPML1 knockdown in MDA-MB-231 cells resulted in significant metabolic disruptions affecting critical pathways such as the tricarboxylic acid (TCA) cycle, amino acid metabolism, nucleotides biosynthesis, and carnitine metabolism (**Figure 8 A**). Within the TCA cycle, ML1-KD MDA-MB-231 cells showed a marked reduction in oxoglutaric acid (also known as alpha-ketoglutarate; α-KG) levels **(Figure 8B)**, a key TCA cycle intermediate with crucial roles in mitochondrial energy metabolism, redox balance, and biosynthetic pathways (*65, 66*). Concomitantly, we observed an increase in 2-hydroxyglutarate (2-HG) **(Figure 8B)**, a metabolite derived from α-KG under conditions of oxidative stress (*67*). The reduction of α-KG likely disrupts nitrogen handling and anabolic synthesis, including lipid and amino acid production (*68*). Interestingly, this metabolic imbalance appears to trigger compensatory increases in other TCA intermediates, such as isocitrate, succinate, and malate, indicating a potential metabolic rewiring that may attempt to maintain TCA cycle flux and support residual mitochondrial function in response to α-KG depletion **(Figure 8B)**.

**Figure 8.**
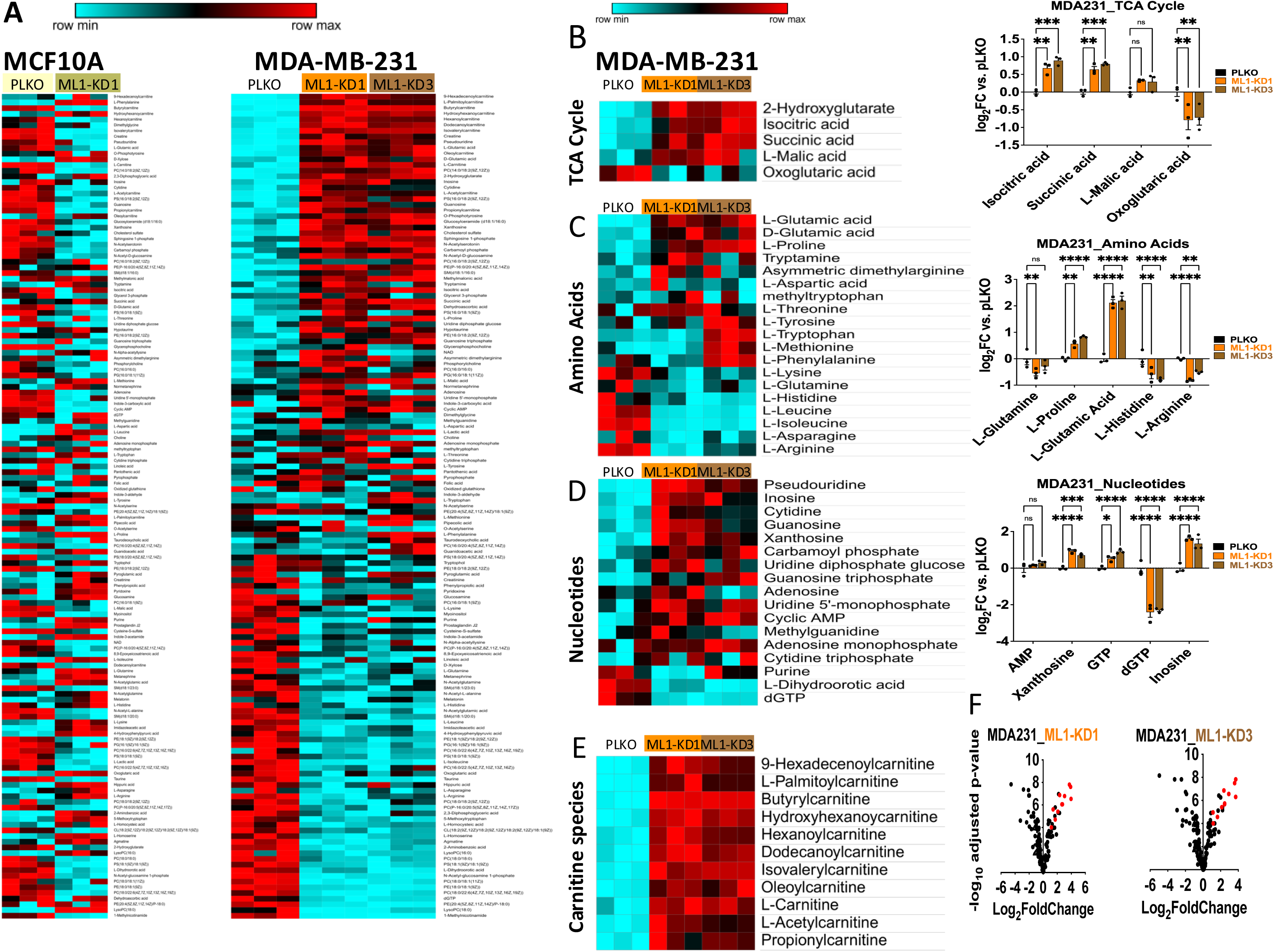
Downregulation of TRPML1 reprograms mitochondrial metabolism and disrupts biosynthetic pathways in MDA-MB-231 Cells. Targeted mass spectrometry-based metabolomic profiling was performed on MDA-MB-231 (TNBC) and MCF10A (non-tumorigenic) cells under control (PLKO) and TRPML1 knockdown conditions (ML1-KD1 and ML1-KD3). (A) Heatmap shows global metabolic alterations upon TRPML1 knockdown, with minimal changes in MCF10A cells and broad remodeling in MDA-MB-231 cells. (B–D) Heatmaps Bar graphs (Mean ± SEM) highlight key changes in metabolites involved in the TCA cycle (B), amino acid metabolism (C), and nucleotide metabolism (D) in MDA-MB-231 cells. ML1-KD increased isocitric acid and succinic acid, with a decrease in oxoglutaric acid, indicating disrupted TCA cycle flux. Amino acid changes included decreased L-glutamine, L-histidine, and L-arginine, and increased L-proline and L-glutamic acid, suggestive of stress-related metabolic shifts and impaired proliferation. Nucleotide alterations included increased xanthosine, inosine, and GMP, with reduced dGTP, pointing to imbalanced nucleotide pools linked to replication stress and cell cycle arrest. (E) Heatmap of carnitine species reveals consistent upregulation in ML1-KD cells, suggesting a incomplete β-oxidation, and a potential shift toward mitochondrial fatty acid metabolism. (F) Volcano plots of ML1-KD1 and ML1-KD3 MDA-MB-231 cells highlight carnitine species as the most significantly upregulated metabolites (dots marked in red), reinforcing the impact of TRPML1 downregulation on mitochondrial substrate handling. Data acquired from a triplicate from one biological experiment. Statistics: one-way ANOVA with Tukey’s multiple comparison test for PLKO vs ML1-KD1/ML1-KD3 of MDA-MB-231. \**P* < 0.05, \*\**P* < 0.01, and \*\*\**P* < 0.001. ns, not significant.

ML1-KD MDA-MB-231 cells also displayed significant reductions in amino acid metabolism, particularly in L-glutamine, histidine and arginine levels **(Figure 8C)**. Glutamine dependence is a metabolic trait of cancer cells, serving as both a nitrogen and carbon source for the TCA cycle, nucleotide biosynthesis, and redox balance (*69, 70*). Its reduction in ML1-KD cells likely compromises these essential functions, further exacerbating metabolic stress. Arginine, which is partly derived from glutamine metabolism (*71*), also declined in ML1-KD cells **(Figure 8C)**. As a precursor for polyamine synthesis and proteins synthesis essential for cancer cell growth (*72–74*), arginine deficiency likely contributes to reduced cell proliferation in ML1-KD cells. The interconnection between arginine and glutamine metabolism highlights the broader metabolic stress imposed by ML1-KD. Interestingly, ML1-KD MDA-MB-231 cells displayed increased proline content **(Figure 8C)**, consistent with previous studies showing proline accumulation in detached cancer cells (*75*). This response has been linked to decreased proliferation and slower cell cycle G1 progression (*75, 76*). Given that ML1-KD cells exhibit mitochondrial dysfunction and growth-arrest phenotype, the elevated proline levels may reflect a similar detachment-like metabolic adaptation.

In addition to disruptions in the TCA cycle and amino acids, TRPML1 knockdown altered purine nucleotide metabolism. Specifically, ML1-KD cells displayed lower levels of dGTP alongside accumulation of xanthosine and inosine **(Figure 8D)**, indicating impaired de novo guanylate synthesis and a compensatory shift toward the salvage pathway (*77, 78*). Guanylates are vital for nucleic acid synthesis and cancer cell survival and their depletion has been associated with induction of cell cycle arrest (*79*) and caspase-independent apoptosis (*80, 81*), consistent with the phenotype observed in ML1-KD cells.

Finaly, our metabolomic data showed that the most upregulated metabolites in ML1-KD cells were carnitines species **(Figure 8E)**. Carnitine facilitates fatty acid transport into mitochondria for energy production but given the impaired mitochondrial function in ML1-KD cells, this likely reflects a failed compensatory response (*82, 83*). Indeed, prior studies showed that elevated carnitines such as C3DC, C4, C10:1, and C10:2 are negatively associated with breast cancer survival (*84*), consistent with the metabolomic stress observed in ML1-KD cells. Altogether, the impact of ML1-KD on mitochondrial function and structure translates into profound metabolic disruptions, underscoring its pivotal role in maintaining mitochondrial integrity and cellular energy homeostasis. These data define a cancer-selective metabolic rewiring downstream of TRPML1, sustaining bioenergetic and biosynthetic programs in TNBC cells.

### 8. TRPML1 Downregulation Enhances the Efficacy of Doxorubicin and Paclitaxel

We next explored whether targeting TRPML1 could enhance the therapeutic potential of chemotherapeutic agents. Because of the high toxicity and dose-limiting side effects of doxorubicin (DOXO) and paclitaxel (PXL), two widely used drugs in breast cancer treatment (*85*), we hypothesized that TRPML1 downregulation would sensitize MDA-MB-231 cells to these chemotherapeutics, potentially improving their efficacy at lower doses. First, we performed a dose-response analysis of both DOXO and PXL on MDA-MB-231 PLKO control cells. We identified 3 nM as a subeffective concentration for both agents that did not significantly alter proliferation (**Figure 9A**). Next, we assessed whether ML1-KD MDA-MB-231 cells would exhibit heightened sensitivity to this suboptimal dose. As shown in **Figure 9B**, at 3 nM, both DOXO and PXL significantly reduced the proliferation of ML1-KD MDA-MB-231 cells, whereas this concentration had no impact on PLKO control cells (**Figure 9B**). These results suggest that TRPML1 knockdown increases the sensitivity of TNBC cells to DOXO and PXL, supporting the potential for effective lower-dose treatment in vitro. Taken together, these findings indicate that loss of TRPML1 creates a chemosensitized state in TNBC cells, highlighting TRPML1 as a tractable target to widen the therapeutic window of standard chemotherapy

**Figure 9:**
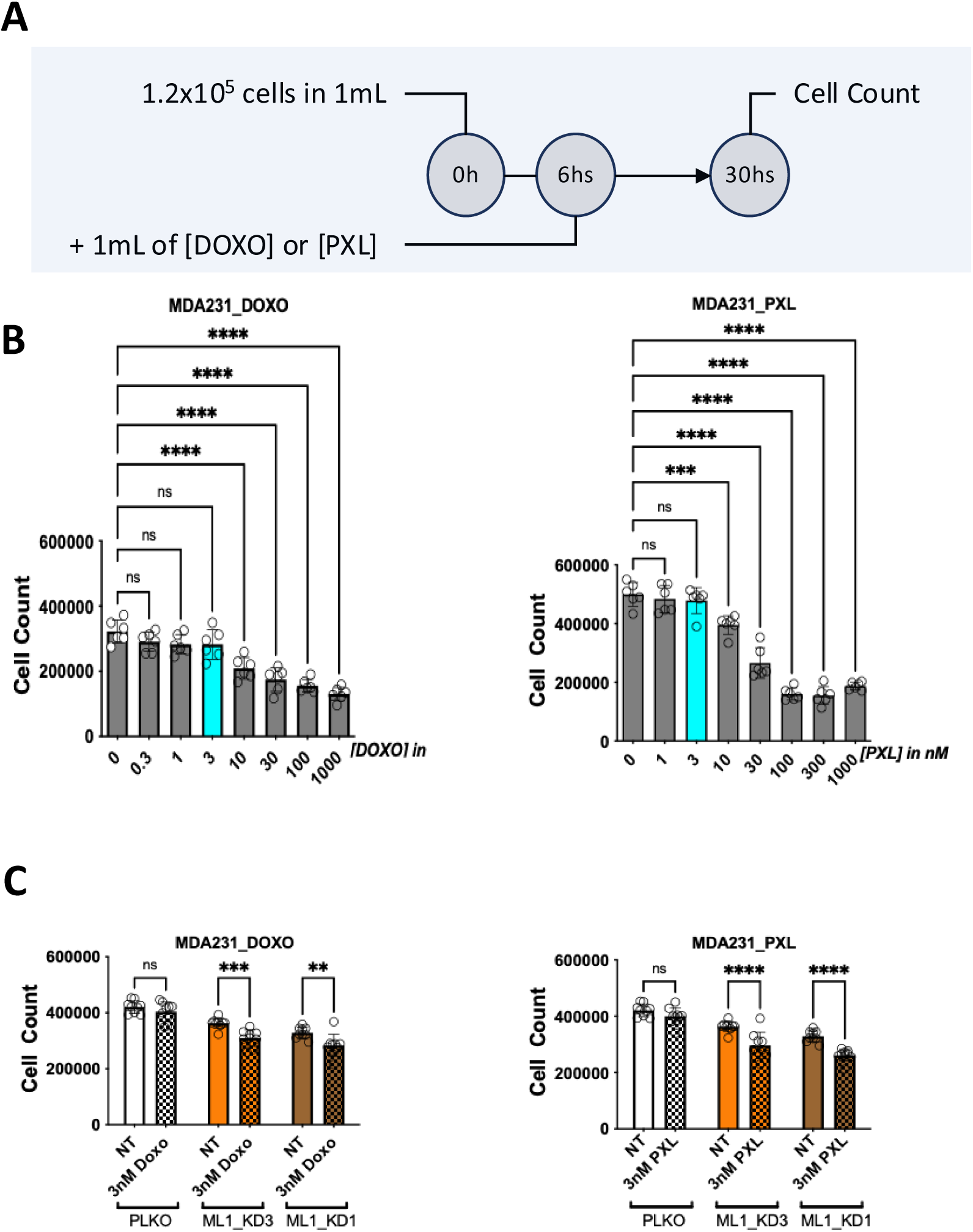
Downregulation of TRPML1 enhances the efficacy of doxorubicin (DOXO) and paclitaxel (PXL) in MDA-MB231 TNBC. (A) schematic presentation of the used protocol for cell treatment and time of counting cells. (A) Dose-response of DOXO and PXL, identifying 3nM as the lowest dose having no effect on MDA-MB-231 PLKO cells, for both DOXO and PXL. (C) ML1-KD re-sensitize MDA-MB-231 to 3nM DOXO and PXL. Data presented in this figure are acquired from 3 independent biological experiment, each with at least six technical repetitions. All graphs (Mean ± SD) show raw cell counting, from 3 independent biological experiment, each between six to nice technical repetitions. Statistics: one-way ANOVA with Tukey’s multiple comparison test for PLKO vs ML1-KD1/ML1-KD3 of MDA-MB-231. \**P* < 0.05, \*\**P* < 0.01, and \*\*\**P* < 0.001. ns, not significant.

## DISCUSSION

Our findings highlight the critical role of the lysosomal Ca^2+^ channel TRPML1 in maintaining lysosomes-mitochondria communication, a relationship that is essential for the metabolic adaptability and survival of TNBC cells. We demonstrate that TRPML1 knockdown disrupts this interplay, leading to mitochondrial dysfunction, impaired autophagy and mitophagy, and altered organelle contact sites, with mitochondria becoming more associated with the ER and less with lysosomes. These changes are cancer-selective (sparing MCF10A) and compromise the cells’ ability to manage metabolic stress and expose vulnerabilities that can be exploited therapeutically. Ultimately, ML1-KD leads to G0/G1 cell cycle arrest, caspase-independent apoptosis, and increased sensitivity to doxorubicin and paclitaxel, suggesting that targeting TRPML1 could offer a novel strategy for enhancing the efficacy of existing TNBC therapies. Together, the data define a TRPML1-dependent program that is non-redundant with other mucolipins in TNBC.

Mechanistically, ML1-KD disrupts several pathways critical for cancer survival (**Figure 10**). (1) It causes lysosomal alkalinization, which impairs both lysosomal degradation and autophagosome-lysosomal fusion, blocking autophagy flux and leading to the accumulation of undegraded cargo (*49–51*). (2) ML1-KD inhibits mTORC1-mediated phosphorylation of ULK1 at Ser757 and activates AMPK-mediated phosphorylation at SerS555, thereby promoting autophagy initiation (*86, 87*). However, the resulting defect in lysosomal acidification prevents completion of the autophagic program, leading to the accumulation of damaged organelles, particularly mitochondria. (3) ML1-KD remodels organellar contact sites, enhancing mitochondria-ER contacts at the expense of mitochondria-lysosomes contacts. Under our conditions, TRPML1 does not directly deliver lysosomal Ca²⁺ to mitochondria: in Ca²⁺-free media, acute TRPML1 agonism fails to raise mitochondrial Ca²⁺ even with TRPML1 overexpression, whereas in the presence of extracellular Ca²⁺, ML1-KD augments cytosolic and mitochondrial Ca²⁺ without changes in MCU abundance or mitochondrial membrane potential—consistent with enhanced plasma-membrane Ca²⁺ entry relayed at reinforced ER–mitochondria interfaces. (4) ML1-KD inhibits TFEB nuclear translocation, downregulating the expression of key genes involved in mitophagy (Pink1 and Parkin) (*17*), mitochondrial biogenesis (TFAM and PGC1α) (*88*), and antioxidant defense (NRF2) (*89*). As a result, mitochondria in ML1-KD cells are less maintained, with increased fragmentation, reduced volume, and lower bioenergetics.

**Fig. 10.**
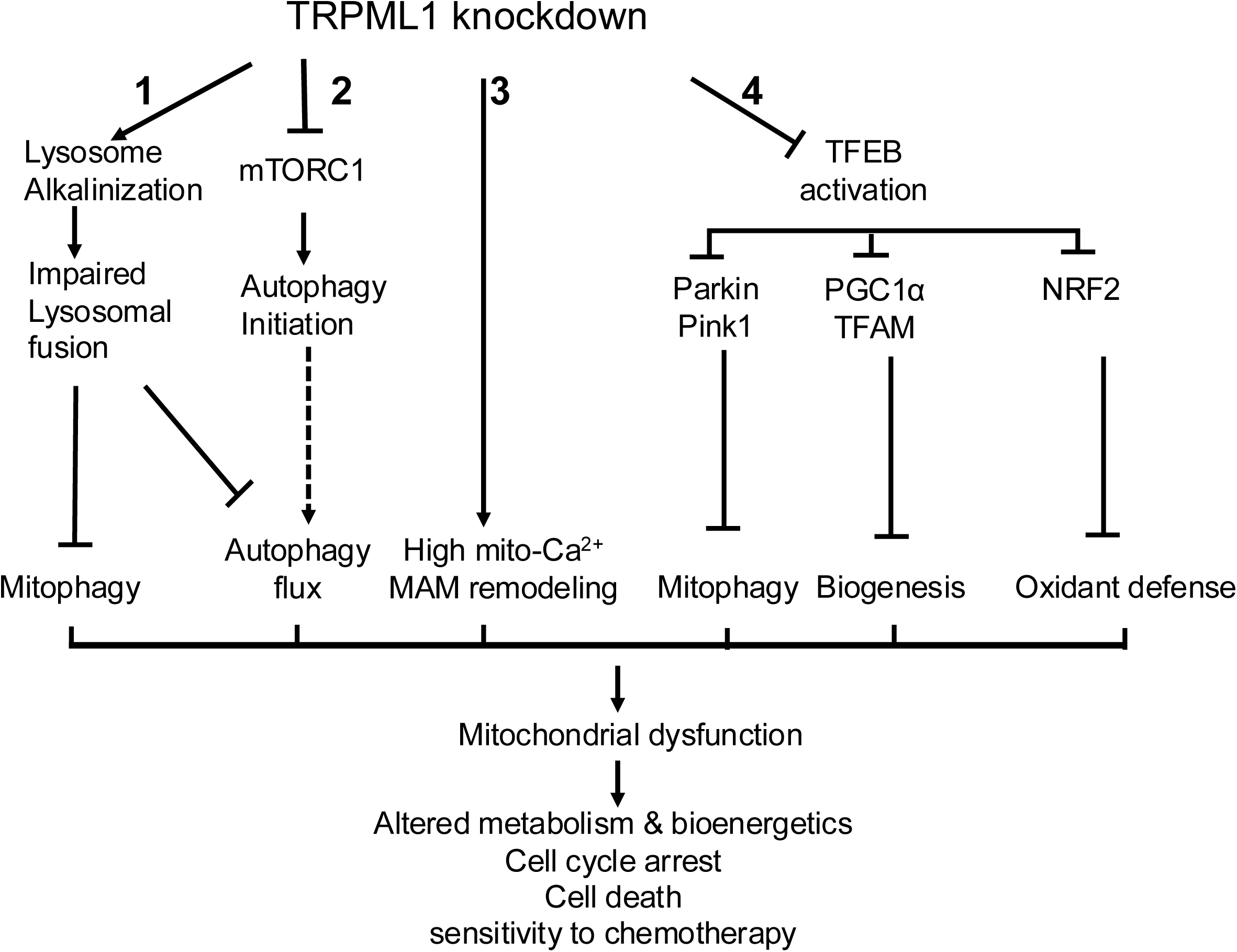
Effect of TRPML1 suppression in TNBC. TRPML1 knockdown (TRPML1 KD) affects three major signaling pathways: 1. Lysosomal Alkalinization and Fusion Impairment: ML1-KD leads to lysosomal alkalinization, increasing lysosomal pH. This impairs the ability of lysosomes to digest cargo and fuse with autophagosomes, thereby blocking proper autophagic flux (Site 1). 2. Autophagy Inhibition via mTORC1: ML1-KD inhibits mTORC1, which initially triggers autophagy initiation. However, due to impaired lysosomal fusion (Site 1), the maturation and termination of autophagy are blocked, leading to defective autophagic flux (Site 2). 3. TRPML1-KD shifts organelle contacts towards less mitochondria/lysosome against more mitochondria/ER crosstalk, thereby elevated mitochondrial Ca2+ uptake. 4. TRPML1-KD-Mediated TFEB Inhibition: ML1-KD disrupts lysosomal Ca^2+^ release, which inhibits calmodulin signaling and results in the accumulation of phosphorylated TFEB. Phosphorylated TFEB is sequestered in the cytosol, preventing its nuclear translocation. Consequently, the downstream genes regulated by TFEB—such as those involved in mitophagy (Pink1 and Parkin), mitochondrial biogenesis (TFAM and PGC1α), and antioxidant defense (NRF2)—are suppressed (Site 4). Disruptions of these signaling culminate in the accumulation of dysfunctional mitochondria due to impaired mitophagy. The mitochondria become fragmented and smaller, altering mitochondrial metabolism and bioenergetics. Ultimately, these disruptions lead to G0/G1 cell cycle arrest, cell death, and increased sensitivity to chemotherapeutic agents.

An important consequence of TRPML1 knockdown is the remodeling of organelle contact sites, with increased mitochondria-ER contacts and decreased mitochondria-lysosomes interactions (#3 in **Fig. 10**). In healthy cells, mitochondria maintain balanced interactions with both the ER and lysosomes, which are essential for efficient Ca^2+^ signaling, cell metabolism, and apoptosis (*14, 21*); whereas in cancer cells these contacts are often dysregulated to support rapid proliferation and metabolic adaptation (*2, 15*). Indeed, lysosomes-mitochondria contacts have been shown to be crucial for mitophagy and mitochondrial quality control (*21, 90, 91*). In ML1-KD TNBC cells, fragmented mitochondria become sequestered within the ER network, distancing them from lysosomes and disturbing their functional coordination. This topology helps explain mitochondrial Ca^2+^ overload, further exacerbating mitochondrial dysfunction and metabolic stress.

The combined effects of these disruptions – lysosomal alkalinization, defective autophagy, TFEB inhibition, altered organelle contacts, and indirect mitochondrial Ca^2+^ overload – create a metabolic crisis in ML1-KD cells. This crisis manifests as significant and profound alterations in metabolic pathways, including the TCA cycle, amino acid metabolism, and nucleotide biosynthesis. Reduced a-KG levels impairs nitrogen balance and the biosynthesis of essential macromolecules (*6, 92*), while decreased glutamine disrupts glutaminolysis, a metabolic pathway often exploited by cancer cells to fuel rapid proliferation (*69, 70*), further exacerbating metabolic stress. Interestingly, our metabolomic profiling revealed a significant accumulation of carnitine species in ML1-KD cells. Carnitine is critical for transporting long-chain fatty acids into mitochondria for beta-oxidation (FAO), a pathway often upregulated in cancer cells under metabolic stress (*93–95*). However, in our model, mitochondrial dysfunction caused by TRPML1 knockdown likely impairs FAO capacity. As a result, the observed increase in carnitine species may not reflect a functional shift toward lipid oxidation but rather a metabolic bottleneck, where some fatty acid transport persists despite a loss of oxidative capacity. Alternatively, it could suggest a disruption in lipid handling or altered mitochondrial import of fatty acids. Future studies will be required to determine the direct mechanistic link between TRPML1 and carnitine metabolism, whether it contributes to the observed vulnerabilities in ML1-KD cells or represents a broader feature of mitochondrial stress response. These metabolic alterations ultimately reduce ATP production, limit the availability of substrates for DNA and protein synthesis, and increase oxidative stress, creating an environment that is incompatible with sustained cell proliferation (*3, 4*). The accumulation of damaged mitochondria due to impaired mitophagy and autophagy flux amplifies oxidative stress and triggers mitochondrial-dependent apoptosis (*34, 96*). The suppression of TFEB-mediated mitochondrial biogenesis and antioxidant pathways further exacerbates the cells inability to cope with metabolic and oxidative stress, providing a mechanistic basis for cell cycle arrest in the G0/G1 phase and caspase-independent apoptosis (*56, 97*).

Importantly, we show that targeting TRPML1 sensitizes TNBC cells to low doses of doxorubicin and paclitaxel. This sensitization likely arises from the convergence of mitochondrial dysfunction, autophagy impairment, and redox imbalance, which weaken the cells’ ability to withstand the cytotoxic effects of chemotherapy (*98, 99*). By disturbing the lysosomal-mitochondrial axis, TRPML1 inhibition creates a metabolic weakness that exacerbates the stress induced by chemotherapeutic agents, leading to enhanced cell death. This approach could potentially leverage the metabolic vulnerabilities of TNBC cells to reduce the adverse side effects associated with high-dose chemotherapy, offering a significant clinical advantage and improving the therapeutic window for TNBC treatment. Notably, whereas TRPML3 targeting in TNBC also enhanced doxorubicin sensitivity through a caspase-dependent route (*31*), the chemosensitization we report here for TRPML1 arises from a distinct cascade, culminating in caspase-independent death. Indeed, although TRPML1 and TRPML3 share a modular architecture and lysosomal localization, they operate in distinct pH windows: TRPML1 conducts at acidic lysosomal pH (*100, 101*), whereas TRPML3 is proton-inhibited and becomes permissive as the lumen approaches neutral/alkaline pH (*102, 103*). This pH gating partitions labor—TRPML1 predominates in normally acidified lysosomes, while TRPML3 emerges when lysosomes neutralize/alkalinize. Functionally in TNBC, (i) death programs diverge: TRPML1 knockdown triggers EndoG-associated, caspase-independent apoptosis, whereas TRPML3 targeting activates caspase-3/7–dependent apoptosis; and (ii) the mTOR/autophagy control nodes differ: TRPML1 knockdown suppresses mTOR, increases initiation but blocks flux at the fusion/acidification step, and dampens TFEB-PINK1/Parkin/PGC-1α/TFAM/NRF2, whereas in our TRPML3 study disrupted lysosomal acidification associated with mTOR activation and mitochondrial dysfunction, with no detectable change in TFEB nuclear translocation (El Hiani laboratory). Importantly, although ML1-KD elevates lysosomal pH, the TRPML1-specific TFEB/mitophagy suppression and contact-geometry remodeling persist, indicating that TRPML3 does not compensate under these conditions. Together, these distinctions indicate that TRPML1 and TRPML3 fine-tune lysosomal outputs through complementary and non-overlapping mechanisms.

Despite these findings, our study has some limitations. While we demonstrated that TRPML1 knockdown selectively affects TNBC MDA-MB-231 cells without impacting non-cancerous MCF10A cells, TNBC is a highly heterogeneous disease of a diverse cellular population. Interestingly, MDA-MB-231 cells themselves exhibit intra-line heterogeneity, making them a representative model for studying this complexity. Our previous *in vivo* data further support the therapeutic potential of TRPML1 inhibition, as ML1-KD significantly suppresses tumor growth and completely prevented lymph node formation in SCID mice (*27*). Nevertheless, it remains to be determined whether TRPML1 inhibition would elicit similar effects in other TNBC cell lines with different genetic backgrounds. Additionally, while we observed no direct effect of TRPML1-mediated lysosomal Ca²⁺ release on mitochondrial Ca²⁺ uptake, the precise molecular mechanisms linking TRPML1 knockdown to enhanced extracellular Ca²⁺ influx and increased mitochondrial Ca^2+^ uptake remain unclear. Future studies should explore these mechanisms in greater details, as they could provide further insights into the role of TRPML1 in Ca^2+^ signaling and its potential as a therapeutic target.

In conclusion, our study underscores the critical role of TRPML1 in maintaining lysosomal and mitochondrial homeostasis in TNBC cells, two organelles whose interdependence is essential for the survival and proliferation of TNBC cells. By targeting TRPML1, we exploit metabolic vulnerabilities that enhance the efficacy of conventional chemotherapies. Thus, TRPML1 represents a promising therapeutic target for developing novel anti-cancer strategies, particularly in aggressive breast cancers such as TNBC. Taken together, our findings support a cooperative—but non-redundant—mucolipin model in which TRPML1 and TRPML3 jointly fine-tune lysosomal signaling, organelle crosstalk, and therapeutic vulnerability in TNBC. Future studies will determine the potential additive effects of targeting both TRPML1 and TRPML3 for enhancing the chemosensitivity of TNBC cells.

## MATERIALS AND METHODS

### Study design

The main objectives of this study were to 1) Elucidate the mechanism by which TRPML1 mediated TNBC cell death, 2) Investigate the impact of TRPML1 on the interplay between lysosomal and mitochondria, and 3) Evaluate the therapeutic potential of TRPML1 inhibition in combinations with chemotherapeutics. To achieve these objectives, we employed a combination of genetic, biochemical, and metabolomic approaches. ML1-KD was achieved using ShRNA against TRPML1 in TNBC cells (MDA-MB-231) and non-cancerous mammary epithelial cells (MCF10A) as a control. Early passages (between 3 and 7) of these cell lines were used to ensure consistency and reproducibility. All experiments were conducted in at least three biologically independent experiments to ensure rigor. Data collection and analysis were performed in a blinded manner whenever possible to minimize bias.

### Cell Culture

MDA-MB-231 (MDA231) and MCF10A were kindly provided by Dr. Shashi Gujar (Dalhousie University), and HEK293T cells were generously provided by Dr. Paul Linsdell (Dalhousie University). All cells are regularly tested for mycoplasma contamination. MDA-MB-231 and HEK293T cells were cultured in Dulbecco’s Modified Eagle Medium (DMEM) supplemented with 10% fetal bovine serum (FBS). MCF-10A cells were maintained in DMEM/F-12 medium containing 5% horse serum, 10 µg/mL human insulin, 20 ng/mL human epidermal growth factor (hEGF), and 0.5 µg/mL hydrocortisone.

### Lentivirus production, stable transduction, and transient infection

Short hairpin RNA (shRNA) clones targeting TRPML1 shTRPML1 were purchased from Dharmacon. Empty plasmid (PLKO) and two different shTRPML1 plasmids (ML1-KD1 and ML1-KD3) were used according to the protocol for the 3rd generation lentiviral packaging system (*104*). Briefly, lentiviral particles were generated in HEK293T cells by co-transfection of PsPAX2 (6 μg) and PMD2G (3 μg), with either the PLKO or ML1-KD (6 μg) plasmids using lipofectamine 2000 (Thermo scientific). The lentivirus was harvested at 48 h post-transfection, filtered (Millex-GS; 0.45-μm sterile filter), and stored at −80 °C. For transduction, MDA231 and MCF-10A cells were seeded in 6-well plates and cultured for 24 h. Medium containing 200 μl of lentivirus and 8 μg/ml of polybrene (Sigma) was added to the cells and allowed to incubate for 48 h, then selected by Puromycin (2 µg/mL) for 4 days. Knockdown efficiency was assessed by RT-qPCR. For this, total RNA was extracted using TRIzol reagent (Invitrogen) and purified with the DNA-free™ DNA Removal Kit (Thermo Fisher Scientific). Complementary DNA (cDNA) was synthesized from 2 µg RNA using iScript Reverse Transcriptase (Bio-Rad). Quantitative PCR (qPCR) was performed using SYBR Green Mater Mix (Bio-Ras) and gene-specific primers (kindly see table 1 for sequences). For transient infection, cells were transfected with RFP-GFP-LC3 or TFEB-GFP or Mito-GFP, and or Lamp1-GFP plasmids using Lipofectamine 2000 (Thermo Fisher Scientific), and imaged or analyzed 24-48 hours post-transfection.

### Cell Proliferation and colony formation assay

For cell proliferation assay, cells were seeded at a density of 5× 10^5^ cells/well and incubated for 72 h at 37 °C/5% CO_2_. After incubation, cells were trypsinised and counted using automated cell counter (CellDrop BF, DeNovix. For the colony formation assay, 500 cells were seeded per well in 6-well plates, and cultured for 14 days with changes media every 72h. On day 14, the media was removed, and cells were gently washed with PBS. Colonies were fixed with 1mL of Coomassie Brilliant Blue (R-250, #1610400, Bio-Rad) dissolved in methanol and Acetic Acid, for 20 min at RT. Stained plates were rinsed by running water for 3 min and allowed to air-dry. Colony images were captured with a phone camera, and the colony formation was evaluated based upon the number of colonies in a 6-well plate using Image J software. For drug tolerance assay, PLKO control and TRPML1 KDs cells, were seeded in 35mm culture dishes at a density of 1.2 × 10⁵ cells per dish in complete DMEM. Cells were allowed to adhere for 6 hours at 37°C in a humidified incubator with 5% CO₂. After attachment, cells were treated with either Doxorubicin or Paclitaxel. Following 24 hours of drug exposure, cells were harvested using trypsinization, and total cell numbers were determined using automated cell counter.

### Western Blot Analysis

PLKO and ML1-KD of MDA231 and MCF10A cells were lysed in radioimmunoprecipitation assay (RIPA) buffer supplemented with protease and phosphatase inhibitors (Thermofisher Scientific). Protein concentrations were determined using the bicinchoninic acid (BCA) assay (Thermofisher Scientific). Equal amounts of protein (1–4 mg/mL) were mixed with 4× Laemmli buffer, boiled, and resolved on SDS-PAGE gels. Proteins were transferred to nitorcellulose membranes and blocked for 1 h in PBS containing 0.1% Tween-20 and 5%BSA. Membranes were incubated overnight at 4C with the following primary antibodies (all from Cell Signaling Technology, 1:1000 dilution unless otherwise noted): Cell cycle regulators: anti-Cyclin B1 (#4138, anti-Cyclin D1(#2922), anti-p27 (#3686), anti-E2F1 (#3742); Apoptosis markers: anti-Cleaved-Caspase 9(#7237), anti-Caspase 9 (#9502), anti-Cleaved-Caspase7 (#9491), anti-Caspase7 (#9492), anti-Cleaved-Caspase3 (9661), anti-Caspase3 (#9662), anti-p-Bcl-2(S70) (#2827), anti-EndoG(#4969), anti-AIF(#4642), Mitochondrial Dynamics: anti-MFN1 (#14739), anti-MFN2 (#11925), anti-OPA1 (#80471), anti-p-DRP1 (S616) (#3455), anti-p-DRP1 (S637) (#4867), anti-MFF (#86668), Mitochondrial bioenergetics: OXPHOS (#ab110411, Abcam), MCU (#14997), Autophagy regulator: anti-p-P70S6K (Thr389) (#9205, Cell Signaling), anti-P70S6K (#9202, Cell Signaling), anti-p-ULK1 (S757) (#6888, Cell Signaling), anti-p-ULK1 (S555) (#5869, Cell Signaling), anti-ULK1 (#8054, Cell Signaling), anti-LC3 A/B (#12741, Cell Signaling), anti-p-AMPK (Thr 172) (#2531, Cell Signaling), anti-AMPK (#2532, Cell Signaling), Mitophagy Regulators, PINK1 (#6946, Cell Signaling), Parkin (#4211, Cell Signaling), Stress Response: anti-Nrf2 (#12721, Cell Signaling), anti-TFAM (#8076, Cell Signaling) anti-PGC1-alpha (#2178, Cell Signaling). After washing, membranes were incubated with IRDye secondary antibodies for 1 hour at room temperature (800CW Goat anti-Rabbit [1:10,000; LicorBio, # 926-32211; Ex/Em: 778/795 nm] or 680RD Goat anti-Mouse [1:10,000; LicorBio, # 926-68070, Ex/Em: 679/696]). Fluorescent signals were visualized using the Odyssey CLx Imager (Licor) imaging system with appropriate infrared fluorescence channels. Band intensity was quantified using Image J software.

### Mitochondrial structure

To assess mitochondrial morphology, cells were seeded at a density of 1 × 10⁵ cells per well onto 18-mm glass coverslips (thickness 0.16–0.19 mm; Mercedes Scientific #R0018)) placed in 12-well plates and allowed to adhere for 48 hours. On the day of imaging, cells were incubated with 150 nM MitoTracker™ CMXRos (Thermo Fisher Scientific, #M7512) in serum free DMEM for 30 minutes at 37°C to label metabolically active mitochondria. Following staining, cells were washed three times with Hanks’ Balanced Salt Solution (HBSS) to remove unbound dye and maintained in HBSS supplemented with 10% FBS during imaging. Live-cell imaging was performed using a Zeiss spinning disc 880 confocal microscope equipped with a 100×/1.46 NA glycerol immersion objective and an environmental chamber maintained at 37°C with 5% CO₂. MitoTracker fluorescence was captured using excitation at 579 nm and emission at 599 nm. Z-stack images were acquired to allow 3D reconstruction of mitochondrial networks. Acquired image stacks were analyzed using Imaris software, following the segmentation approach described by Taguchi et al. (2021). Mitochondria were identified and segmented based on individual organelle volume and sphericity. For classification, mitochondrial structures were grouped into three distinct morphological categories: fragmented mitochondria were defined by a sphericity value greater than 0.88, intermediate forms had sphericity values between 0.77 and 0.88, and elongated mitochondria displayed sphericity values below 0.77. The proportion of each category was determined by calculating the total mitochondrial volume in each group relative to the total mitochondrial volume per cell.

### Lysosomal pH

To evaluate lysosomal acidification, cells were prepared under the same conditions as described for mitochondrial structure analysis. After 48 hours of adhesion on 18-mm glass coverslips in 12-well plates, cells were incubated with 150 nM LysoTracker™ Red DND-99 (Thermo Fisher Scientific, #L7528) in complete medium at 37°C for 30 minutes to selectively label acidic organelles. Following staining, cells were washed three times with phosphate-buffered saline (PBS) to remove excess dye and then incubated in fresh complete medium for an additional 30-minute chase period to ensure selective retention of the probe in acidic lysosomal compartments. Live-cell imaging was performed on a Zeiss LSM 880 confocal microscope, and LysoTracker fluorescence was recorded using excitation at 577 nm and emission at 590 nm. High-resolution 2D images were captured under identical acquisition parameters across all samples to allow for comparative quantification. For analysis, individual cells were manually segmented using Fiji (ImageJ) software, and the mean fluorescence intensity of the LysoTracker signal was measured per cell. To ensure accuracy, background fluorescence was subtracted from each measurement prior to analysis. Lysosomal fluorescence intensity was used as a proxy for lysosomal acidification, with lower fluorescence indicating elevated lysosomal pH.

### Intracellular ATP Measurement

Cellular ATP levels were quantified using the Luminescent ATP Detection Assay Kit (Abcam, ab113849) following the manufacturer’s instructions. PLKO control and ML1 knockdown cells were seeded in white-walled, flat-bottom 96-well plates (5,000 cells/well, triplicates) and cultured for 48 hours at 37°C with 5% CO₂. At the time of assay, plates and reagents were equilibrated to room temperature, and media was gently aspirated. Cells were lysed by adding 50 µL of lysis buffer (Detergent) per well and incubated for 5 minutes on a shaker. Meanwhile, the substrate solution was prepared by reconstituting the lyophilized enzyme mix (Substrate) in substrate buffer. Immediately after lysis, 50 µL of the reconstituted substrate solution was added to each well. Plates were covered to protect from light and incubated at room temperature for 15 minutes to allow luminescence stabilization. Luminescence was measured using a Synergy H1 microplate reader (BioTek) with a 1-second integration time. An ATP standard curve (10 µM to 0.01nM) was generated using serial dilutions of the supplied ATP standard. ATP levels were normalized to cell number, quantified from parallel wells using manual cell counting.

### Flow Cytometry Analysis

All flow cytometry assays were performed on a BD FACS Celesta™ cytometer (BD Biosciences) equipped with a 488-nm laser. A minimum of 20,000 events per sample were collected, and data were analyzed using FlowJo software (v10.7.1). Cells were gated based on forward and side scatter to exclude debris and doublets. *Apoptosis* was assessed using Annexin V (BioLegend, #640906) and 7-Aminoactinomycin D (7-AAD) (BioLegend, #420404) staining. Cells were harvested 72 hours post-treatment, washed with Annexin binding buffer (BioLegend, #422201), and incubated with 12.5 μg/mL Annexin V and 20 μg/mL 7-AAD in the dark for 15 minutes at RT. Fluorescence of Annexin V in PLKO and ML1-KD cells was measured by FITC channel, analysed by FlowJo software and presented as %. ***Cell cycle*** distribution was determined using propidium iodide (PI) staining. Cells were synchronized by serum starvation for 12 hours, then released in complete medium for 10 hours to resume cell cycle progression (*31, 105*). Following harvesting and washing, cells were fixed in ice-cold 70% ethanol overnight at 4°C. On the day of analysis, cells were washed with PBS and stained with 100 µg/mL PI and 50 µg/mL RNase A (Sigma) in PBS for 1 hour at RT in the dark. DNA content was analyzed in PI channel, analysed by FlowJo software, and presented as %. *Total ROS* levels were measured using the oxidation-sensitive dye CMH₂DCFDA (Thermo Fisher Scientific, #D399). Cells were incubated with 5 µM CMH₂DCFDA in serum free DMEM for 60 minutes at 37°C, then washed twice with PBS and resuspended in 1 mL PBS for acquisition. Mean fluorescence intensity was measured in the FITC channel. *Mitochondrial ROS* was quantified using MitoSOX™ Red (Thermo Fisher Scientific, #M36008). Cells were incubated with 5 µM MitoSOX in serum free DMEM for 30 minutes at 37°C in the dark. After washing, cells were analyzed immediately, and red fluorescence was measured in the PE channel.

### Autophagosome–Lysosome Fusion Assay

Cells were seeded on 18-mm glass coverslips in 12-well plates and transfected with the tandem fluorescent reporter plasmid RFP-GFP-LC3 (Addgene #21074) using Lipofectamine 2000 (Thermo Fisher Scientific) according to the manufacturer’s protocol. Twenty-four hours post-transfection, cells were washed with PBS and fixed in 4% paraformaldehyde for 15 minutes at RT, then mounted using ProLong™ Diamond Antifade Mountant (Thermo Fisher, #P36961). Fluorescence imaging was performed using a Zeiss LSM 880 confocal microscope equipped with a 63× oil immersion objective. GFP and RFP were sequentially excited at 488 nm and 561 nm, respectively, and emissions were collected at 509 nm (GFP) and 583 nm (RFP). 2D images were acquired using identical acquisition settings across all samples. Images were analyzed using Fiji (ImageJ). RFP⁺/GFP⁺ (yellow) puncta were counted as autophagosomes, while RFP⁺/GFP⁻ (red-only) puncta were counted as autolysosomes. Quantification was performed manually on a minimum of 20 cells per condition, per experiment, and results were plotted as the ratio of yellow to red puncta (*47*).

### TFEB Nuclear Translocation Assay

Cells were seeded on 18-mm glass coverslips and transfected with GFP-NI-TFEB plasmid (Addgene #38119) using Lipofectamine 2000 (Thermo Fisher Scientific). Twenty-four hours post-transfection, cells were fixed with 4% paraformaldehyde for 15 minutes at RT before imaging. Imaging was performed using a Zeiss LSM 880 confocal microscope with a 40× oil immersion objective. GFP was excited at 488 nm and emission was collected at 509 nm. The number of cells with nuclear TFEB was counted and plotted as % of TFEB nucleus positive cells/total cells in the field. At least 10 fields, each 10-15 cells, per condition, per experiments, were analyzed and compared across PLKO, ML1-KD, and ML-SA1–treated and untreated PLKO cells.

### Three-Dimensional Organelle Contact Analysis

To assess organelle contact remodeling, cells were seeded on 18-mm glass coverslips and transfected with Mito-GFP (for mitochondria), LC3-mCherry (for lysosomes), or Sec61-NEON (for Endoplasmic Reticulum) plasmids. After 24 h, cells were stained with MitoTracker CMXRos (for mitochondria) or LysoTracker Red DND-99 (for lysosomes), depending on the studied organellar contacts. After staining, cells were washed three times with HBSS and maintained in HBSS supplemented with 10% FBS during live-cell imaging. Confocal Z-stacks were acquired using a Zeiss LSM-880 microscope equipped with a 100× oil immersion objective and an environmental chamber (37°C, 5% CO₂). Z-step size was set 0.3 μm to ensure Nyquist sampling for volumetric reconstruction. Fluorophores were imaged using sequential acquisition with fixed laser power and detector settings across all samples. We used the Airyscan detector to achieve higher resolution and sensitivity. This allowed us to visualize cellular organelles more clearly and in greater detail compared to conventional confocal imaging (*106*). Image analysis was performed using Imaris 10.2 software. Surfaces were generated for mitochondria, lysosomes, and ER compartments using the “Surfaces” module, with intensity thresholds set based on automated background subtraction followed by manual adjustment to capture organelle boundaries. For colocalization of organelles, surface-surface extension in Imaris software was employed. Briefly, “Surface–Surface Contact Area” tool was applied to calculate the contact volume between organelles within a maximum 3D contact distance of 0.2 μm. This distance was chosen to reflect proximity consistent with potential functional tethering (*107, 108*). Contact parameters were normalized to the total mitochondrial or lysosomal surface volume per cell to account for differences in cell size. A minimum of 10 cells per condition, per experiment, were analyzed. Data were reported as contact volume (μm³) or surface area (μm²) per cell and expressed as relative fold-change versus PLKO controls.

### Mitochondrial and Cytosolic Ca^2+^ imaging

To measure both mitochondrial and cytosolic Ca^2+^, 50,000 cells were seeded onto 25 mm glass coverslips (Chemglass #CLS-1760) in complete media and allowed to adhere overnight. Coverslips were then mounted into Attofluor^TM^ cell chambers (Thermo Fisher Scientific #A7816) and cells were suspended in complete media, to which Ca^2+^ sensing dyes were added. For cytosolic Ca^2+^ measurements, the mounted coverslips were incubated with 2 µM Fura-2 AM (Thermo Fisher Scientific #F1225) for thirty minutes at room temperature. Cells were then washed four times with HEPES Buffered Saline Solution (HBSS; 140 mM NaCl, 4.7 mM KCl, 1.13 mM MgCl_2_, 2.0 mM CaCl_2_, 10 mM glucose, and 10 mM HEPES, pH adjusted to 7.4 with NaOH). The chambers were subsequently mounted on an Olympus IX71 Fluorescence Microscope. Cells were imaged with a 20X Fluor objective, and the Fura-2 was excited at 340 and 380 nm using a CoolLED pE-340^fura^ shutter with emissions captured at a wavelength of 510 nm with a Hamamatsu ORCA-Spark camera. The ratio of the fluorescence at 340 and 380 nm (F_340_/F_380_) was analyzed with HCImage software (https://www.hcimage.com) by drawing regions of interest around the perimeter of approximately 30 cells per coverslip. For mitochondrial Ca^2+^ measurements, the mounted coverslips were incubated with 100 nM MitoTracker^TM^ Green FM (Thermo Fisher Scientific #M7514) and 1 µM Rhod-2 AM (Thermo Fisher Scientific R1244) for thirty minutes at room temperature. Cells were then washed four times with HBSS and mounted on a Leica Stellaris 5 confocal microscope. Cells were imaged with an HC PL APO CS2 20x/0.75 DRY objective, and MitoTracker^TM^ Green fluorescence was excited at 490 nm with emissions captured at 516 nm, while Rhod-2 AM was excited at 552 nm with emissions captured at 581 nm. Using the Leica Application Suite X (LAS X) (https://www.leica-microsystems.com), regions of interest were drawn around the mitochondria of approximately 30 cells per coverslip. MitoTracker^TM^ and Rhod-2 fluorescence were normalized to the fluorescence at t = 0, and the Rhod-2 signal was ratioed against the MitoTracker^TM^ signal to correct for any changes in focus and eliminate any background cytosolic Rhod-2 signal.

### Metabolomics Analysis

Global metabolomics was performed to quantify steady-state intracellular metabolites in control (PLKO) and TRPML1 knockdown (KD) MDA-MB-231 and MCF10A cells. Cells were scraped in cold (−20°C) 80% methanol and centrifuged at 13,000 × g for 5 minutes. A 25 μL aliquot of the resulting supernatant was added to 225 μL of hydrophilic interaction liquid chromatography (HILIC) loading buffer containing 95% acetonitrile, 2 mM ammonium hydroxide, and 2 mM ammonium acetate. After centrifugation at 13,000 × g for 5 minutes, triplicate 50 μL injections of each sample were loaded onto an Acquity UPLC BEH Amide column (1.7 μm, 2.1 × 100 mm; Waters #186004801). Targeted metabolite detection was carried out by multiple reaction monitoring (MRM) using a Sciex 5500 QTRAP mass spectrometer, with parameters and acquisition settings as previously described (Almasi et al., 2020). Peak heights for individual metabolites were extracted using Skyline software (MacCoss Lab Software). Significantly altered metabolites were defined as those showing ≥1.5-fold change between KD and PLKO conditions. Pathway enrichment analysis was performed to identify functional metabolic clusters using publicly available metabolic pathway databases. Data were presented as fold change relative to PLKO, and statistical comparisons were performed using Student’s *t* test on at least three independent biological replicates.

### Statistics

Statistical analyses were conducted using GraphPad Prism 9.0.1 (GraphPad, La Jolla, CA, USA). Data are presented as means ± SD of at least three biologically independent experiments. For comparisons between two groups, an unpaired Student’s t-test was used. For comparisons involving more than two groups, one-way or two-way ANOVA with Tukey’s multiple comparisons to adjust for multiple comparisons. A P-value of less than 0.05 was considered statistically significant. Figure legends specify the statistical tests used for each experiment.

## DATA AVAILABILITY

All data needed to evaluate the conclusions in the paper are present in the paper. Additional data related to this paper may be requested from the authors.

## ACKNOWLEDGMENTS

We thank Dr. G. Gerard, Facility Manager of Cellular & Molecular Digital Imaging Facility at Dalhousie University for assistance with optimization and analysis of confocal Imaging. We thank Dr. Dong for providing the Phospho-P70S6K and Phospho-Ser757ULK antibody in early stages of the project.

## AUTHOR CONTRIBUTIONS

A.K.S. designed experiments and collected data for all figures of the paper. S.A. initiated the project and performed experiments in Fig. 1,2 and assisted with experiments in Fig.9. J.C.B. Performed experiments in Fig.7. B.E.K. performed experiments in Fig.2 and assisted with Fig.8. R.E.Y and SM.E. performed experiments for Fig. 4A-F. L.S. Performed experiments in Fig.8. S.P. performed experiment in Fig.2G. V.V.V. Performed experiments in Fig.2A-E. S.G. Designed and assisted with experiment in Fig.2. T.P. assisted with the conceptualization, design and supervision of the overall study, revised, edited and approved the final version of the manuscript. M.T. and Y.EH. led the conceptualization and design of the study, supervised the research, secured funding, drafted the original version of the manuscript, and revised, edited, and approved the final version.

## Additional Information

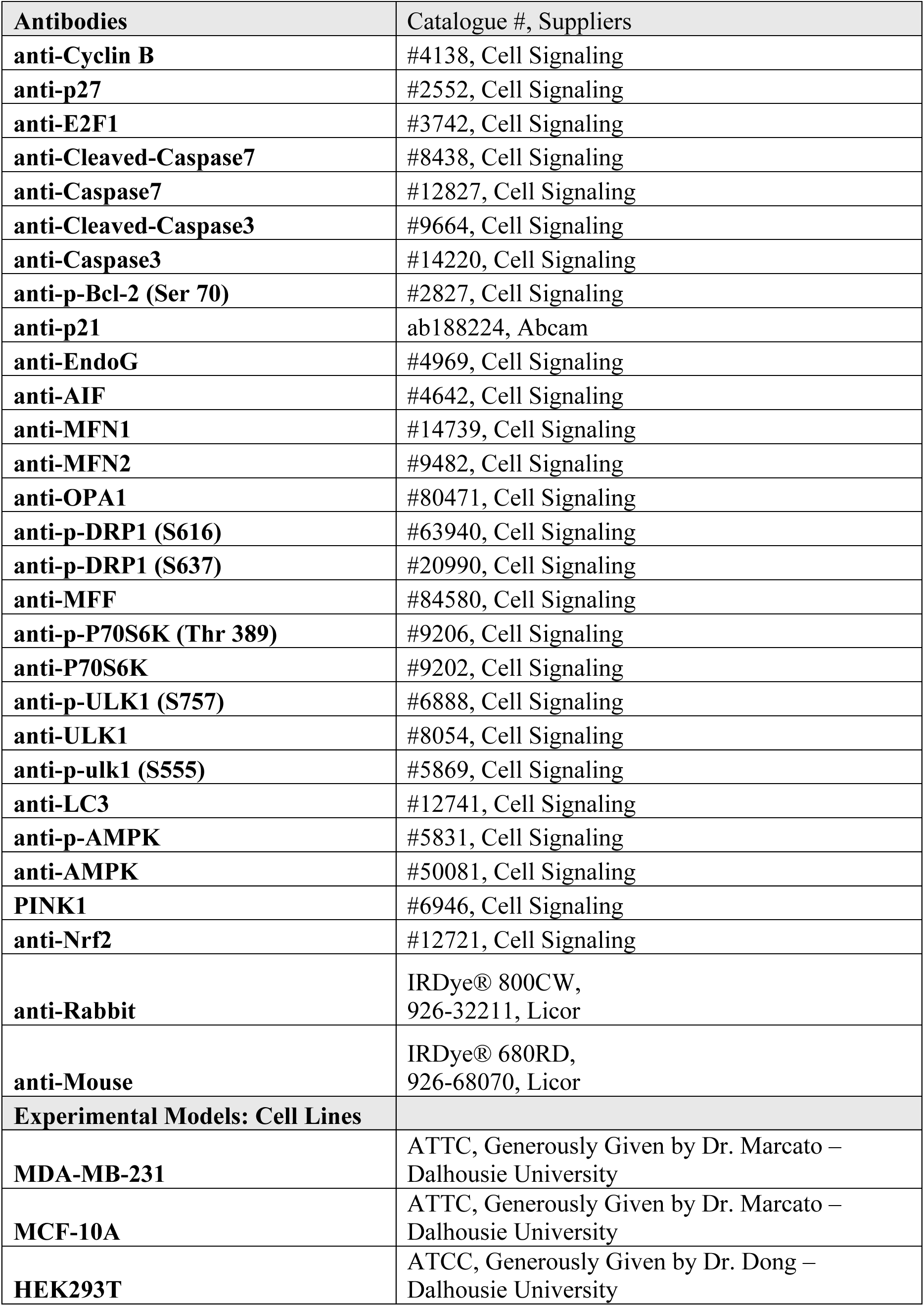
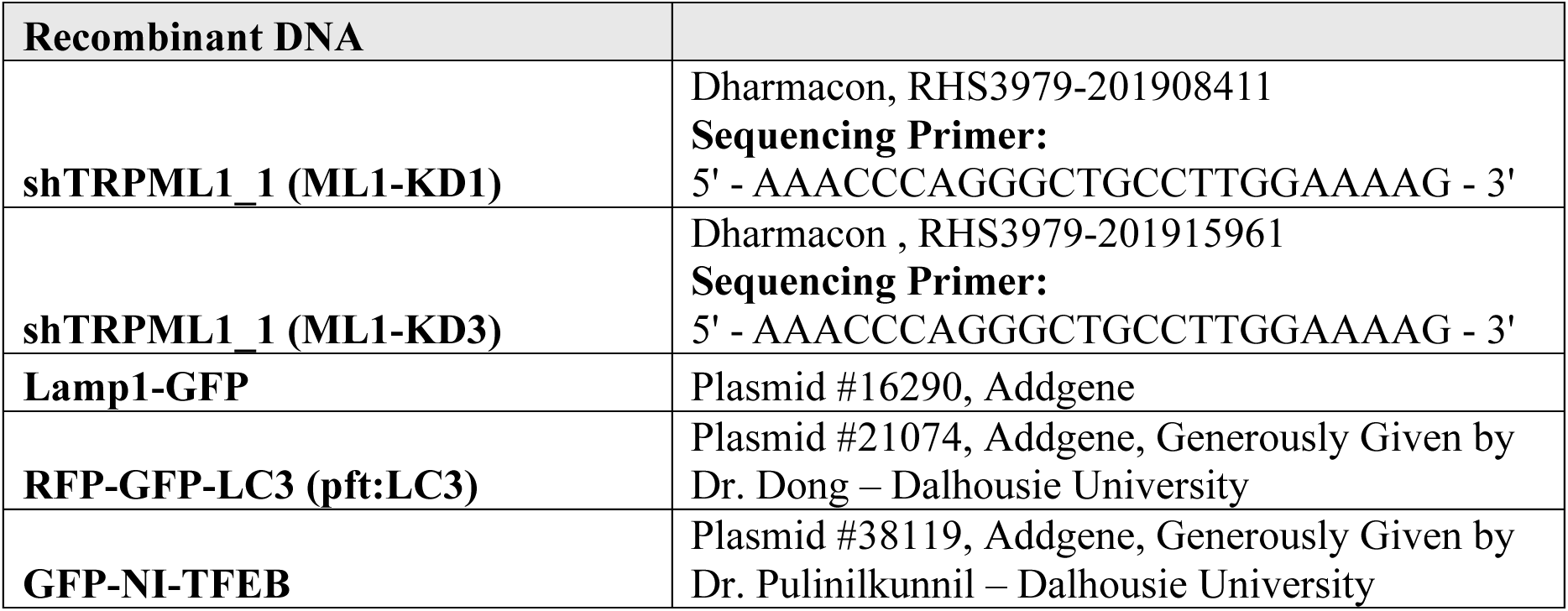

